# Experience-dependent plasticity of gustatory insular cortex circuits and taste preferences

**DOI:** 10.1101/2022.08.16.504203

**Authors:** Hillary C. Schiff, Joshua F. Kogan, Maria Isaac, Lindsey A. Czarnecki, Alfredo Fontanini, Arianna Maffei

**Affiliations:** Dept. of Neurobiology and Behavior SUNY Stony Brook; Stony Brook, NY, USA; Graduate Program in Neuroscience, SUNY Stony Brook; Stony Brook, NY, USA; Medical Scientist Training Program, SUNY Stony Brook; Stony Brook, NY, USA

**Keywords:** insular cortex, taste experience, inhibitory plasticity, sensitive periods, perineuronal nets

## Abstract

Early experience with food influences taste preference in adulthood. How gustatory experience influences development of taste preferences and refinement of cortical circuits has not been investigated. Here we exposed weanling mice to an array of tastants and determined the effects on the preference for sweet in adulthood. We demonstrate an experience-dependent shift in sucrose preference persisting several weeks following the termination of exposure. A shift in sucrose palatability, altered neural responsiveness to sucrose, and inhibitory synaptic plasticity in the gustatory portion of the insular cortex (GC) were also induced. The modulation of sweet preference occurred within a restricted developmental window, but restoration of the capacity for inhibitory plasticity in adult GC reactivated the sensitivity of sucrose preference to taste experience. Our results establish a fundamental link between gustatory experience, sweet-preference, inhibitory plasticity, and cortical circuit function, and highlight the importance of early life experience in setting taste preferences.

## Introduction

Taste preferences guide food choices thereby impacting health and quality of life. At the transition from relying on mother’s milk to foraging, animals learn to independently taste food and act upon it, approaching and consuming nourishing food, or avoiding and rejecting dangerous substances. In animals, preferences for key tastes such as sweet are viewed as innate (Yiannakas and Rosenblum, 2017), yet they can be influenced by early life experience (Galef and Henderson, 1972; Capretta and Rawls, 1974; Mennella et al., 2001; Volcko et al., 2020). Even in human infants, early taste experiences modulate gustatory preferences later in life (Mennella et al., 2001; Ventura and Mennella, 2011).

In adults, the gustatory portion of the insular cortex, the gustatory cortex (GC), plays a central role in a variety of taste-related functions, including integrating information about identity and hedonic value of taste (Katz et al., 2001; Fontanini and Katz, 2006), predicting the occurrence of a taste based on food’s anticipatory cues (Veldhuizen et al., 2011; Samuelsen et al., 2012; Gardner and Fontanini, 2014; Livneh et al., 2017), learning about safety or danger related to tastes (Berman and Dudai, 2001; Haley et al., 2016; Schier et al., 2019), forming associations between tastes and postingestive effects (de Araujo et al., 2008; Oliveira-Maia et al., 2012), and making taste-informed decisions (Parkes and Balleine, 2013; Schiff et al., 2018; Vincis et al., 2020). These findings highlight GC plasticity and point to the possibility that circuits in GC may be sensitive to experience with taste. While the role of experience in shaping preferences has been demonstrated in in flies (May et al., 2019; Vaziri et al., 2020), there is currently no information about the relationship of taste experience during postnatal development, taste preference, and neural circuit function in gustatory cortical circuits.

Here we investigated whether taste experience at weaning modulates the preference for sweet later in life, and assessed the neural underpinnings of taste exposure in mouse GC. We report that early life exposure to tastes has profound and persistent effects on sweet preference, neural responses to sucrose, and the postnatal maturation of inhibitory circuits in GC. The same exposure started in adulthood did not affect sucrose preference, indicating the presence of a sensitive period for the experience-dependent modulation of sweet preference. Manipulation of GC inhibition in adult mice was sufficient to reopen the sensitive window and restore taste-dependent plasticity in adult mice. To our knowledge, this is the first study to directly link early life dietary experience with cortical circuit plasticity and responsiveness to tastants, indicating a powerful effect of the integration of chemosensory experience and nutrition on brain development.

## Results

### Taste exposure early in life enhanced preference for sucrose

We developed a taste exposure paradigm to manipulate taste experience (Fig. 1). Weanlings were given repeated access to 4 different solutions (“early exposure”, EE; Fig. 1a) in their home cage drinking bottle on an 8-day schedule (Fig. 1a; 150mM sucrose (S) on days 1 and 5, 20mM citric acid (CA) on days 2 and 6, Ensure (E) on days 3 and 7, and 100mM salt (NaCl) on days 4 and 8). Following exposure, EE mice returned to a diet of chow and water until the end of the experiment. A separate group of littermates was assigned to the control group (“naïve”) and maintained on a standard diet of chow and water. In separate sets of control experiments, a group of naïve mice was used to confirm that all solutions were consumed comparably or more than water. Sucrose and Ensure were preferred over water, consistent with an innate preference for sweet taste, while NaCl and citric acid were consumed comparably to water (Supplementary Fig. 1a). We also verified that there was no difference in body weight between mice in the naïve and EE groups during or after EE (Supplementary Fig. 1b).

**Figure 1.**
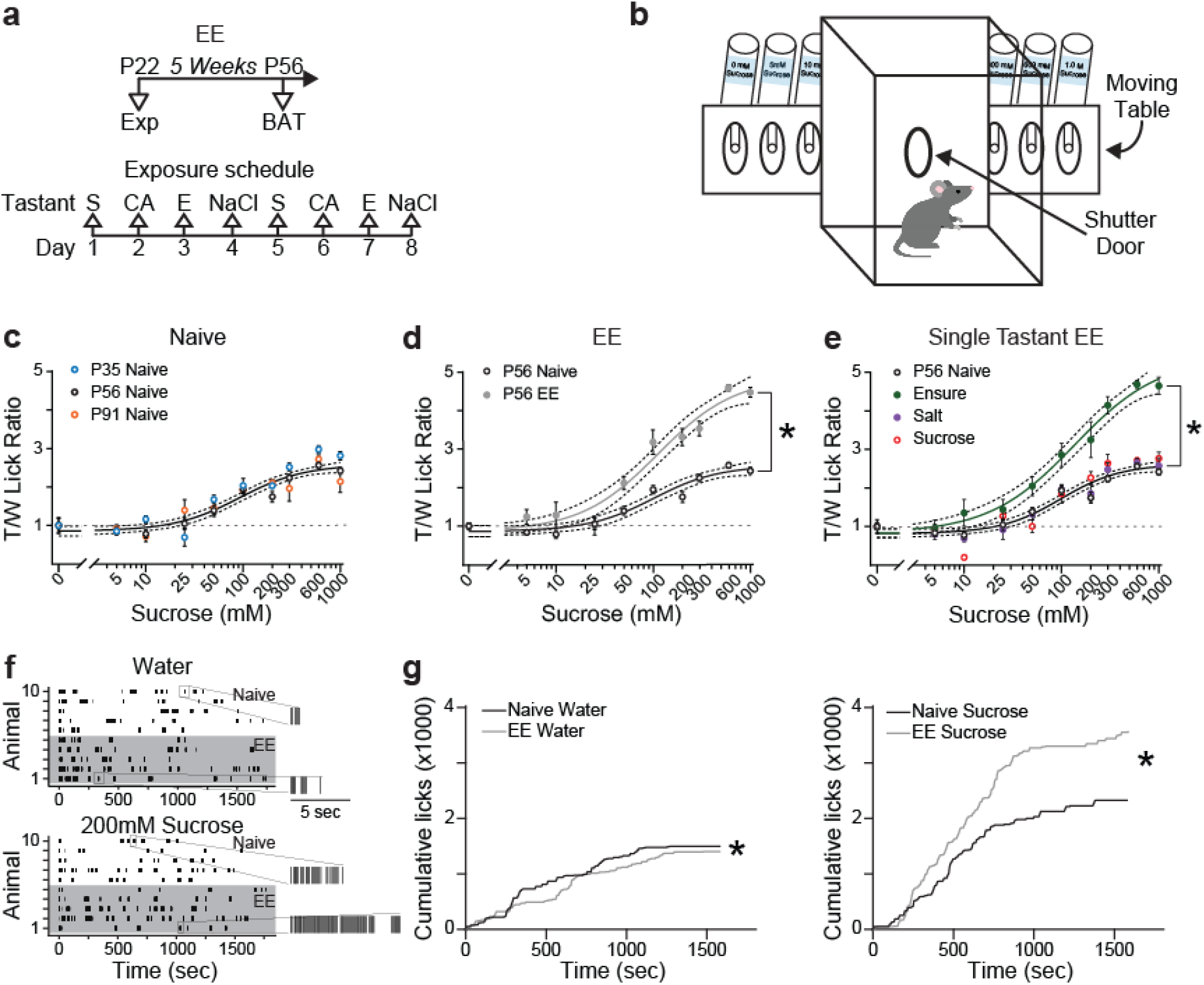
Early Exposure (EE) induces long-lasting changes in sucrose preference. **a.** Top: schedule for EE and testing. Bottom: sequence of taste solutions. **b.** Representation of Davis rig gustometer. **c.** Sucrose curves from naïve mice by age group. A single curve fits all datasets (p=0.071, F test). **d.** Sucrose curve for P56 naïve (black) and EE (grey) mice. Separate curves were needed to fit the datasets (p<0.001, F test). **e.** Sucrose curves for P56 mice in modified EE. Separate curves were needed for Ensure (p<0.001, F test), while a single curve was sufficient to fit naïve (black), NaCl (purple), and sucrose (red) (p=0.115, F test). For c-e: Dark line: curve fit; dotted lines: 95% confidence intervals. Dotted grey line plotted at 1 shows response to water (0mM sucrose). Data points: Mean±sem. **f-g.** Analysis of licking for P55-P58 naïve and EE mice tested on a 30-min trial of access to water or 200mM sucrose. **f.** Lick raster plots for water (top) and sucrose (bottom). Shaded area: licks by mice in the EE group. **g.** Left: accumulation of licks to water from naïve (black) and EE (grey) mice (p=8.2×10^-10^, Mann-Whitney U test). Right: accumulation of licks to sucrose in naïve (black) and EE (grey) mice (p=5.2×10^-5^, Mann-Whitney U test). Asterisks indicate p ≤ 0.05.

We used a brief access test (BAT) to determine the effects of EE on sweet preference. Preference was assessed by measuring licking to different concentrations of sucrose with a Davis rig gustometer (Smith et al., 1992; Smith, 2001) (Fig. 1b) and quantifying the tastant-to-water (T/W) lick ratio for each stimulus concentration. BAT exposes mice to different concentrations of sucrose (in mM: 0, 5, 10, 25, 50, 100, 200, 300, 600, 1000) for multiple, brief (10 seconds) trials. There were no developmental changes in sucrose preference in distinct cohorts of naïve mice from 3 distinct age groups (Fig. 1c; P35 n=7, P56 n=26, P91 n=10 mice, F_(6,420)_=1.956, p=0.071, F test), suggesting that sucrose preference is stable in mice maintained on their regular diet, as previously shown (Inui-Yamamoto et al., 2017). In contrast, sucrose preference was enhanced in mice from the EE group when assessed either at P35 (Supplementary Fig. 1c) or at P56 (Fig. 1d), one or four weeks from the end of EE, respectively. Separate sigmoid functions were required to fit the data sets from age matched naïve and EE mice (Fig. 1d; P56: naïve n=26, EE n=7 mice; F_(3,323)_=79.76, p<0.0001, F test), and the slope of the sucrose preference curves were significantly different (Supplementary Fig. 1d). Unless otherwise stated, our analysis of preference focused on P56 in order to examine the long-term effects of taste experience. For each experimental condition, a group of naïve and one of EE littermates were run in parallel.

The observed shift in sucrose preference did not depend on familiarity with sucrose, as 8 days of exposure to sucrose alone did not shift the preference curve measured at P56 (Fig. 1e, Supplementary Fig. 1d), in line with previous observations (Wurtman and Wurtman, 1979). Experience with salt may also influence sweet preference (Smriga et al., 2002), but 8 days of exposure to NaCl alone did not affect the sucrose curve (Fig. 1e, Supplementary Fig. 1d; naïve n=26, sucrose EE n=7, salt EE n=8 mice; F_(6,399)_=1.720, p=0.115, F test;). However, 8 days of Ensure induced a shift comparable to that observed with the EE paradigm (Fig. 1e; naïve n=26, Ensure EE n=7 mice; F_(3,318)_=83.06, p<0.0001, F test;), and increased the slope of the sucrose curve (Supplementary Fig. 1d).

To determine whether the shift in sucrose curve represents increased palatability for this tastant, we analyzed licking microstructure, a widely used approach to assess different aspects of taste perception (Spector et al., 1998). We compared licking behavior in naïve and EE mice which had access to water or 200mM sucrose for a 30-minute trial on subsequent days (Fig. 1f). The number of bouts of licks for water and sucrose was comparable (Supplementary Fig. 2a), but in EE mice bout size (measured as licks/bout) for sucrose was significantly larger (Supplementary Fig. 2b). Finally, mice in the EE group accumulated licks for water more slowly than naïve mice (Fig. 1g, left), while they accumulated licks more rapidly for sucrose (Fig. 1g, right; water: naïve n=1496 licks, EE n=1405 licks, U=927887, p=8.2×10^-10^; sucrose: naïve n=2324 licks, EE n=3583 licks, U=3872876, p=5.2×10^-5^, Mann-Whitney U test). As bout size is directly linked to palatability (Davis et al., 1999; Johnson, 2018; Volcko et al., 2020), and accumulation of licks is evidence of increased avidity to sucrose (Roebber et al., 2015), these results indicate that mice in the EE group perceived sucrose as more palatable than naïve mice.

To assess whether the shift in the sucrose curve represents shift in preference specific to sucrose or it extends to other tastants, we compared the curve for salt and saccharin from separate groups of naïve and EE mice. EE did not change the preference for salt (Supplementary Fig. 3a). However, the preference for saccharin was significantly increased in mice from the EE group (Supplementary Fig. 3b), suggesting that experience-dependent changes in sucrose preference generalize to other sweet tastes but do not extend to salt.

The shift in sucrose preference observed in the EE paradigm and in the Ensure EE (Fig. 1e) could have resulted from the variety of factors, such as exposure to tastants and nutrients, postingestive effects due to caloric content, and/or the olfactory component of Ensure. To begin testing the role of postingestive effects, we ran a calorie-free version of EE in which sucrose was replaced with saccharin, and Ensure was replaced with monosodium glutamate (MSG). This paradigm preserves the oral sensation of taste and variety of taste qualities (sweet, sour, salt, umami), but does not engage postingestive mechanisms. The calorie-free exposure did not shift the sucrose preference curve (Fig. 2a; naïve n=26, calorie-free EE n=9 mice; F_(3,343)_=0.7430, p=0.527, F test) nor the slope of the curve (Fig 2a, right; t_(58)_=0.458, p>0.999, Bonferroni-corrected t-test), indicating that calories play a role in the experience-dependent modulation of sucrose preference. To further assess the importance of the postingestive contributions, we tested a cohort of mice which received 8 days of Ensure by tube-feeding beginning on P22. The control group received water in the same manner. While this version of EE exposed mice to calories and engaged postingestive mechanisms, it occurred in the absence of the oral sensation of taste. We found that exposure to Ensure bypassing the mouth did not affect the sucrose preference curve (Fig. 2b; water n=5, Ensure n=7 mice; F_(3,113)_=1.936, p=0.128, F test) nor its slope (Fig 2b, right; t_(10)_=0.8322, p=0.425, unpaired two-tailed t-test). These results point at integration of orosensory and enteric information in the experience-dependent modulation of sweet preference.

**Figure 2.**
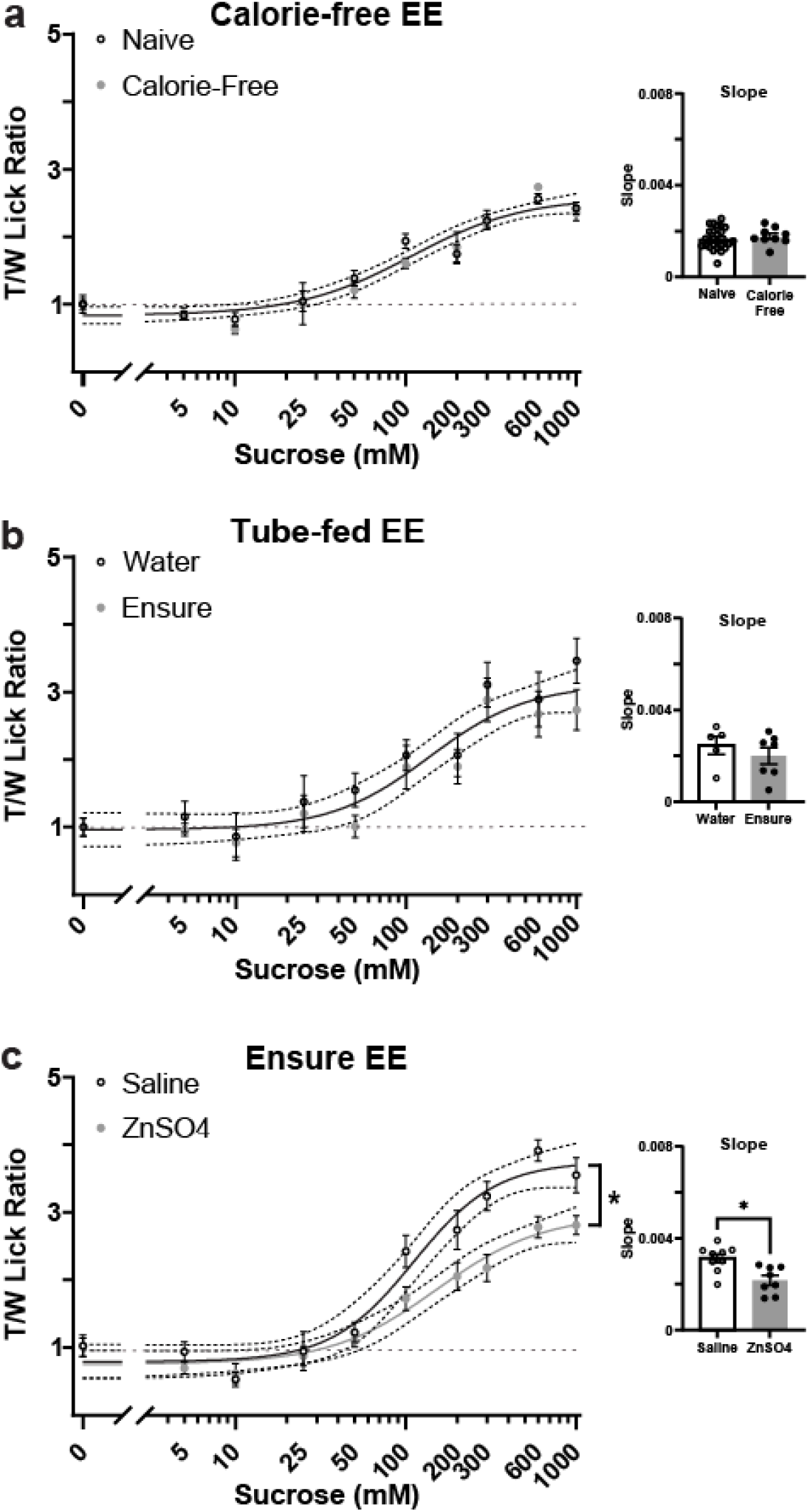
EE depends on integration of orosensory, postingestive and olfactory activity. **a.** Sucrose curve for P56 naïve (black) and calorie-free EE (grey) mice. A single curve was sufficient to fit both datasets (p=0.527, F test) and the slope of the curves did not differ (p>0.999, t-test with Bonferroni correction for multiple comparisons). **b.** Sucrose curve for P56 mice tube-fed with water (black) or with Ensure (grey) from P22-P29. A single curve was sufficient to fit both datasets (p=0.128, F test) and the slope of the curves did not differ (p=0.425, t-test). **c.** Sucrose curve for P56 mice with Ensure EE treated with either saline or ZnSO_4_ at P21. Separate curves were needed to fit the datasets (p=6.349×10^-11^, F test) and the slope of the preference curve were significantly different (p=0.0038, t-test). Note that the sucrose curve and the slope of the curve in ZnSO_4_ mice indicate decreased preference. Asterisks indicate p ≤ 0.05.

To assess the possible contribution of the olfactory component of Ensure, we induced temporary anosmia during exposure (Alberts and Galef, 1971; Slotnick and Gutman, 1977). Mice received an intranasal instillation of either saline (control group) or ZnSO_4_ (temporary anosmia group) prior to receiving Ensure EE. This manipulation did not affect the amount of Ensure consumed (Supplementary Fig. 4a; saline n=9, ZnSO_4_ n=8 mice; F_(1,15)_=0.07741, p=0.785, 2-way RM ANOVA main effect of drug), nor body weight (Supplementary Fig. 4b; p=0.207, ANOVA main effect of drug). We verified anosmia and recovery of smell in a separate group of mice (Supplementary Fig. 4c, d; see also Supplementary Fig. 4 legend for details). Mice rendered temporarily anosmic during Ensure EE showed a reduction in the sucrose preference curve compared to intact Ensure-exposed mice (Fig. 2c; saline n=9, ZnSO_4_ n=8, F_(3,163)_=19.65, p=6.349×10^-11^, F test) accompanied by a decrease in the slope of the preference curve (Fig 2Fc, right; saline n=9, ZnSO_4_ n=8 mice, t_(15)_=3.420, p=0.0038, two-tailed unpaired t-test), indicating that the olfactory component of Ensure contributed to the experience-dependent enhancement in sucrose preference. Together, the results of these assays support the conclusion that early experience with taste solutions led to a persistent increase in sucrose preference and palatability, and that the modulation of sucrose preference depends on the integration of sensory (gustatory and olfactory) and postingestive components.

### Experience-dependent modulation of neural responses to sucrose

Motivated by its role in taste processing and taste guided behaviors (Kusumoto-Yoshida et al., 2015; Vincis and Fontanini, 2016; Yiannakas and Rosenblum, 2017; Vincis et al., 2020), we focused on the gustatory insular cortex (GC) as a possible circuit regulating taste preferences. We used two-photon calcium imaging in awake naïve and EE mice to measure the sucrose-evoked responses of GC neurons during active licking for various concentrations (Fig. 3a, b; in mM: 0, 25, 50, 100, 200, 600). We used the offspring of PV-cre (Taniguchi et al., 2011) and floxed-tdTomato mice (Madisen et al., 2010) (Fig. 3a, c) in order to record calcium signals simultaneously from pyramidal and parvalbumin-positive (PV+) neurons. We, extracted ΔF/f traces, and deconvolved activity using a widely adopted algorithm (see methods, Fig. 3d-e) (Pnevmatikakis and Giovannucci, 2017). We focused on PV^+^ GABAergic neurons because they are highly sensitive to changes in postnatal sensory experience in other cortices (Fagiolini et al., 2004; Hensch, 2005; Di Cristo et al., 2007; Rupert and Shea, 2022) and influence taste-based memory formation in GC (Yiannakas et al., 2021). We recorded 5224 neurons over 12 sessions from 5 mice for the naïve group (4994 non-PV neurons, putative pyramidal, PYR; 230 PV^+^ neurons), and 4795 neurons in 15 sessions from 5 mice for the EE group (4573 PYR; 222 PV^+^ neurons). Neural activity was monitored during the baseline period of the trial (2-second window before trial onset). Following EE, this baseline activity was reduced in PYR neurons (Fig. 3f, g top; naïve n=4994, EE n=4573 neurons, p=0.013, Kolmogorov-Smirnov (K-S) test) while it was increased in PV+ neurons (Fig. 3f, g bottom naïve n=230, EE n=222 neurons, p=7.92×10-9, K-S test), suggesting that the GC circuit leans towards higher inhibition in EE mice. In addition, fewer PYR neurons were activated by sucrose (Fig. 3h; naïve: n=729/4994 (14.6%); EE: n=387/4573 (8.5%); χ^2^(1) = 87.19, p<0.0001, χ^2^ test), while the number of PV^+^ neurons responsive to sucrose was unchanged (Fig. 3h; naïve: n=18/230 (7.8%) vs EE: n=20/222 (9%); χ^2^(1)=0.2053, p=0.69, χ^2^ test). Thus, following EE, the ratio of sucrose-activated excitatory to inhibitory neurons in GC was reduced. To investigate whether the reduced activation of putative PYR neurons modulated their responsiveness to sucrose concentrations, we fitted average responses to each concentration with a linear function and calculated the slope and 95% confidence interval of the fit (see methods, Fig. 3i-j). We observed an increased proportion of neurons with a significant fit in the EE group compared with naïve, and this effect was specific to the sampling window (Fig. 3k, baseline: naïve n=123/729 (16.9%), EE n=68/387 (17.6%), χ^2^(1)=0.087, p=0.77; sampling: naïve n=123/729 (16.9%), EE n=101/387 (26.0%), χ^2^(1)=13.41, p<0.001, χ^2^ test). There was also a significant increase in the absolute slope of the fits (Fig. 3l; baseline: t_(1)_=0.096, p=0.92, sampling: t_(1)_=2.16, p<0.05, unpaired two-tailed t-test). Thus, following EE, the spontaneous activity of PYR neurons decreased, while that of PV neurons increased shifting the E/I balance of the GC network. Consistent with a decreased E/I ratio, fewer PYR neurons responded to sucrose. However, the ones that did respond were more sensitive to sucrose concentrations, indicating that EE had persistent effects on GC circuit excitability, circuit responsiveness and on the coding for sucrose concentrations.

**Figure 3.**
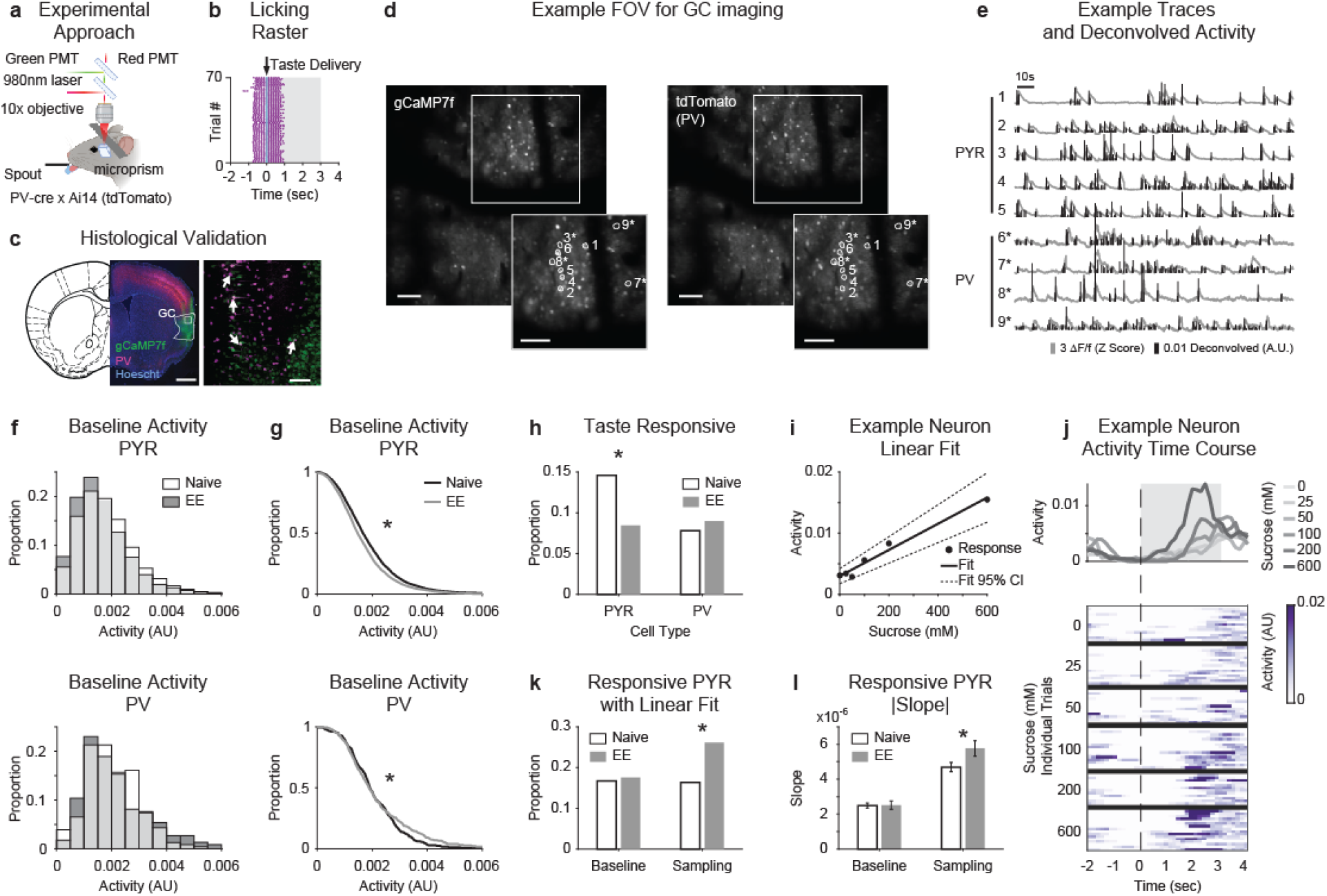
EE induces long-lasting changes in GC neural responses. **a.** Experimental approach for microprism-based calcium imaging in GC. **b.** Representative lick raster showing trial structure, grey shading indicates response window. **c.** Left: representative coronal section showing gCaMP7f expression, PV^+^ neurons, and Hoechst 33342 counterstaining, scale bar=1mm. Right: high magnification view of inset. Arrows: overlapping PV^+^ neurons with gCaMP7f, scale bar=100μm. **d.** Representative two-photon field of view (FOV) for both gCaMP7f (left) and tdTomato (right), scale bar=100μm. Insets show region of interest (ROI) in both channels for 9 example neurons (* indicates tdTomato^+^), scale bar=100μm. **e.** Example traces showing ΔF/f and deconvolved activity for PYR and PV neurons labeled in D. **f.** Distribution of activity at baseline for all PYR (top, naïve: n=4994 EE: n=4573) and PV (bottom, naïve: n=230 EE: n=222) neurons. Naïve in white, EE in dark grey, light grey region represents area where naïve and EE overlap. **g.** Cumulative distribution of activity at baseline for all PYR (top, p=0.013, K-S test) and PV^+^ (bottom, p=7.92×10-9, K-S test) neurons. **h.** Proportion of taste responsive PYR (white bar, naïve: 729/4994; grey bar, EE: 387/4573) and PV+ (white bar, naïve: 18/230, grey bar, EE: 20/222) neurons (PYR: p<0.0001; PV: p=0.65, χ^2^ test). **i.** Representative neuron with a significant linear fit to sucrose response. Black circles indicate response to each sucrose concentration, solid black line shows the linear fit and dashed lines show 95% confidence interval of fit. **j.** Activity from neuron shown in i; dashed line indicates time of taste delivery. Top: Peristimulus time histogram (PSTH) showing average responses to each sucrose concentration. Bottom: heat map showing individual trial responses sorted by concentration (each row represents one trial, 96 trials total). **k.** Proportion of PYR neurons with a significant linear fit in the baseline (white bar, naïve: 123/729; grey bar, EE: 68/387; p=0.77, χ^2^ test) and sampling (white bar, naïve: 123/729; grey bar, EE: 101/387; p<0.001, χ^2^ test) windows. **l.** Absolute slope of PYR neurons with a significant linear fit (same neurons as K, baseline: p=0.92, sampling: p<0.05, unpaired t-test). For panels g, h, k, and l. Asterisks indicate significant difference between naive and EE.

### Taste experience early in life enhances inhibitory synaptic transmission in GC

Reduced spontaneous activity, response suppression in PYR neurons and increased spontaneous activity of PV^+^ neurons may be due to reduced excitability of PYR neurons, increased excitability of PV^+^ neurons, or changes in synaptic transmission. To test these possibilities, we used patch clamp recordings from acute slice preparations obtained from naïve and EE mice at P56 (Fig. 4, Supplementary Fig. 5). For these experiments, as for the imaging *in vivo*, we used the offspring of PV-cre (Taniguchi et al., 2011) and floxed-tdTomato mice (Madisen et al., 2010), to visualize PV^+^ and PV^−^ neurons, and record from both PV and PYR populations. Recorded neurons were filled with biocytin for post-hoc reconstruction of morphology, location, and immunohistochemistry. We confirmed the GABAergic nature of recorded PV^+^ neurons, and the excitatory nature of putative PYR neurons using post hoc immunohistochemistry for GAD67 (Fig. 4a, Supplementary Fig. 5a, e). We obtained input/output functions by injecting square current pulses of different amplitudes and quantifying the frequency of action potentials (Supplementary Fig. 5b, f). Naïve and EE mice showed no differences in either PYR or PV^+^ neurons input/output curves (Supplementary Fig. 5c, g) or input resistance (Supplementary Fig. 5d, h). Thus, EE does not modulate the intrinsic excitability in either PYR or PV^+^ neurons.

**Figure 4.**
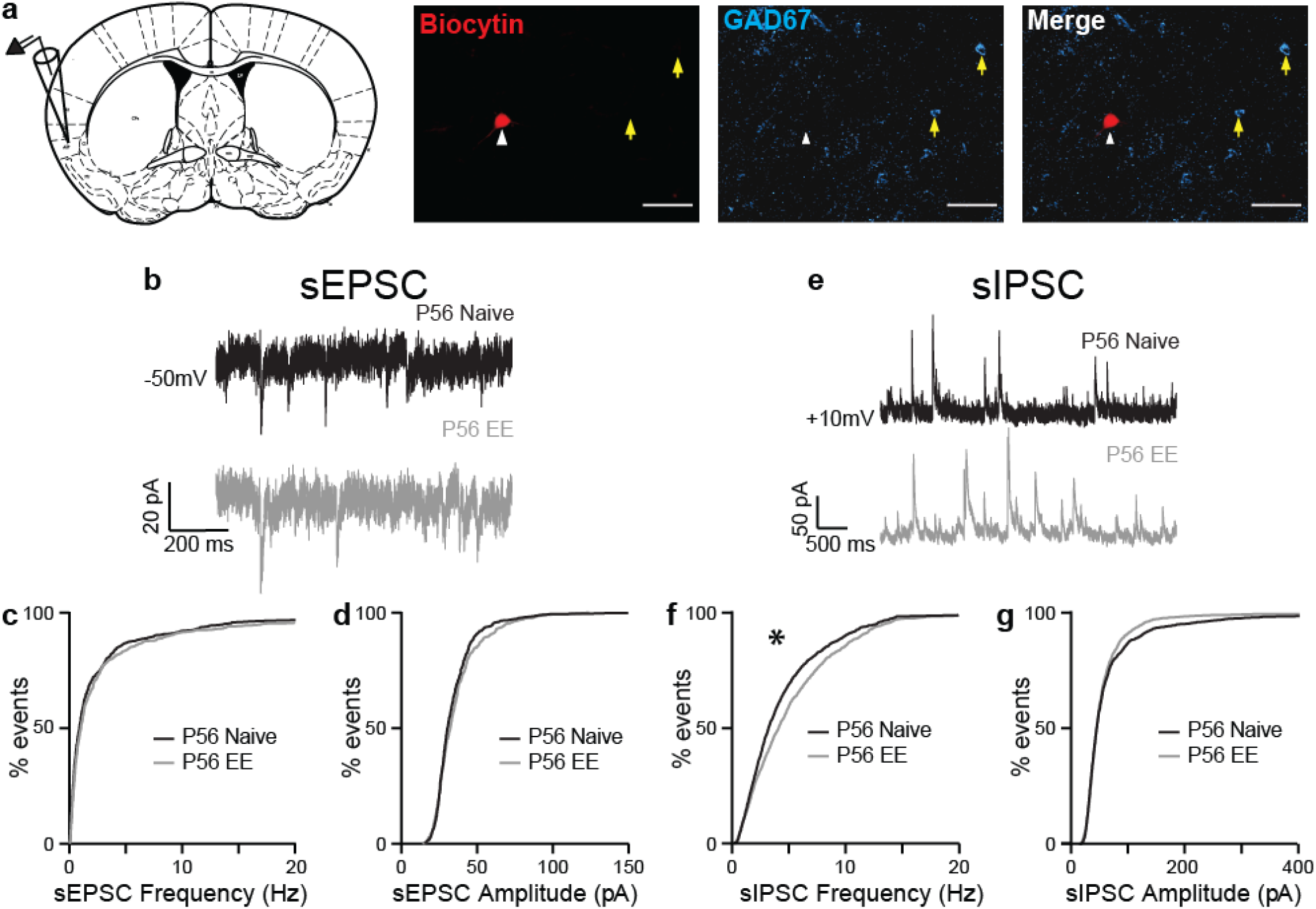
Inhibitory synaptic transmission in GC is modulated by taste experience. **a.** Left: diagram of recording configuration and location of GC. Right: example recorded neuron lacking GAD67 immunostaining (red: biocytin-filled neuron soma imaged through a single z-plane; cyan: GAD67 immunostaining). White arrowhead: recorded neuron; yellow arrows: GAD67-expressing neurons. Scale bar=50μm. **b.** Sample traces of sEPSCs in mice from naïve (black) and EE (grey) groups. Neurons were held at −50mV, the reversal potential for Cl^−^, during recordings. **c.** Cumulative distribution of sEPSC instantaneous frequency (p=0.232, K-S test). **d.** Cumulative distribution of sEPSC amplitude (p=0.455, K-S test). **e.** Sample traces of sIPSCs in mice from naïve (black) and EE (grey) groups. Neurons were held at +10mV, the reversal potential for cations, during recordings. **f.** Cumulative distribution of sIPSC instantaneous frequency (p=9.24×10^-12^, (K-S) test). **g.** Cumulative distribution of sIPSC amplitude (p=0.137, K-S test). Asterisks indicate p ≤ 0.05.

To test the possibility that EE modulates synaptic transmission onto GC PYR neurons, voltage clamp recordings were obtained from confirmed PYR neurons in acute slice preparations of P56 naïve and EE wild-type mice (Fig. 4a, see also Supplementary Fig. 6). Spontaneous excitatory and inhibitory postsynaptic currents (sEPSCs and sIPSCs) were obtained by holding neurons at the reversal potential for chloride (−50mV in our experimental conditions) or for AMPA- and NMDA-mediated currents (+10mV; Fig. 4b, e). Following EE, there were no significant changes in sEPSC amplitude and frequency (Fig. 4b-d; naïve n=15 cells from 10 mice, EE n=14 cells from 11 mice; frequency: D=0.0539, p=0.232; amplitude: D=0.0651, p=0.455, Kolmogorov-Smirnov (K-S) test), while we observed a significant increase in sIPSC frequency, but not amplitude (Fig. 4e-g P56: naïve n=16 cells from 8 mice, EE n=15 cells from 12 mice, sIPSC frequency, D=0.130, p=9.24×10-12, K-S test; sIPSC amplitude, D=0.0416, p=0.137, K-S test). The effect of EE on inhibitory transmission was already detectable at P35 (Supplementary Fig. 6) and persisted at least until P56 (Fig. 4e-g). These results suggest that increased inhibitory synaptic transmission underlies the decreased activation of putative PYR neurons, possibly contributing to modulating sucrose responsiveness and preference.

### Taste experience modulates anatomical markers of inhibitory maturation in GC

To further investigate the effect of EE on inhibitory circuits in GC we quantified anatomical markers, focusing on PV immunofluorescence and the accumulation of perineuronal nets (PNNs), extracellular matrix proteins that preferentially accumulate around PV^+^ neurons (Pizzorusso et al., 2002; Berardi et al., 2004; Sorg et al., 2016). Changes in PV immunofluorescence have been associated with the effects of postnatal experience (Murase et al., 2017) and learning-related plasticity in adult animals (Donato et al., 2013), whereas the accumulation of mature PNNs is associated with the duration of sensitive periods for experience-dependent plasticity (Pizzorusso et al., 2002) and limits the capacity for inhibitory plasticity (Pizzorusso et al., 2006; Gogolla et al., 2009; Sorg et al., 2016). To determine whether EE affected PV immunofluorescence and/or PNN accumulation, we used immunohistochemistry with an antibody against PV and incubation with Wisteria Floribunda Agglutinin (WFA) to label PNNs in wild-type C57BL/6 mice (Fig. 5a). We quantified the intensity of PV and WFA fluorescence from individual PV^+^ neurons and PNNs, as well as the proportion of PV^+^ neurons associated with a PNN in naive and EE mice. In GC of naïve mice, the proportion of PV^+^ relative to all neurons was stable from P35 to P56 (Fig. 5b; P35 n=5, P56 n=4 mice; t_(7)_=0.7685, p=0.467; unpaired two-tailed t-test), as was the PV fluorescence intensity of neurons with and without a PNN (Supplementary Fig. 7a, b). There was also no difference in the proportion of PV^+^ neurons associated with a PNN (Supplementary Fig. 7a, b insets), in WFA fluorescence intensity in P35 and P56 mice (Supplementary Fig. 7c), and in the proportion of PNNs (Supplementary Fig. 7c inset). These data indicate that PV immunofluorescence and accumulation of PNNs of naïve mice reached stable levels in the P35-P56 age group.

**Figure 5.**
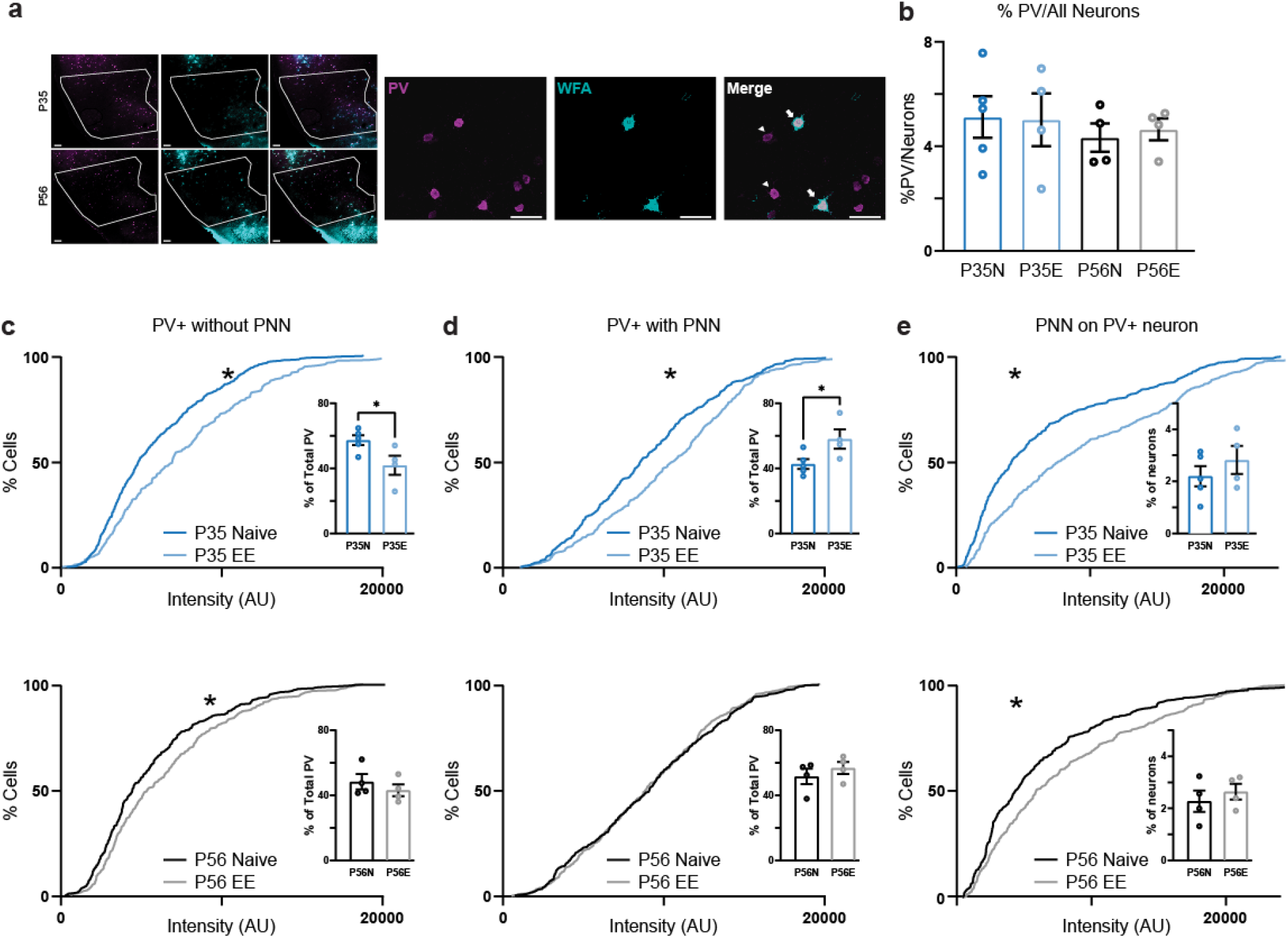
PV^+^ neurons and PNNs associated with PV^+^ neurons are modulated by EE. **a.** Left: example images of GC sections stained with WFA (cyan) in mice from naïve and EE groups. Scale bar=100μm. Right: Sample images of PV immunostaining (left, magenta), WFA staining for PNNs (middle, cyan), and merged view (right). Arrows show examples of double-labeled neurons (PV + PNN) while arrowheads show examples of PV^+^ neurons without a PNN. Scale bar=50μm. **b.** The proportion of PV^+^ neurons in GC was not different between P35 and P56 (p=0.467, t-test), nor was it affected by EE at either age (P35 naïve vs P35 EE p=0.939, t-test; P56 naïve vs P56 EE p=0.661, t-test). **c.** Cumulative distribution of PV immunofluorescence intensity for PV^+^ neurons without a PNN and bar plot showing the proportion of PV^+^ neurons lacking a PNN. Top: at P35, PV immunofluorescence signal increased following EE (p=7.99×10^-6^, K-S test), while the proportion of PV^+^ neurons lacking a PNN was reduced (inset). Bottom: at P56, PV immunofluorescence signal was increased following EE (p=0.016, K-S test) while the proportion of PV^+^ neurons lacking a PNN remained unchanged by EE (inset). **d.** Cumulative distribution of PV immunofluorescence intensity for PV^+^ neurons with a PNN and bar plot of proportion of PV^+^ neurons with a PNN. Top: at P35, PV immunofluorescence signal was increased following EE (p=2.467×10^-5^, K-S test), as was the proportion of PV^+^ neurons with a PNN (p=0.042, t-test). Bottom: at P56, PV immunofluorescence of PV+ neurons with a PNN was unaffected by EE (p=0.745, K-S test), and the proportion of PV^+^ neurons with a PNN remained stable (p=0.416, t-test). **e.** Cumulative distribution of WFA fluorescence intensity for PNNs associated with PV^+^ neurons and proportion of PNNs associated with PV^+^ neurons in GC. Top: at P35, WFA fluorescence increased following EE (p=2.555×10^-8^, K-S test). The proportion of PNNs in GC was not changed by EE (p=0.363, t-test). Bottom: at P56, WFA fluorescence increased following EE (p=8.869×10^-4^, K-S test). The proportion of PNNs was not changed by EE (p=0.497, t-test). Asterisks indicate p ≤ 0.05.

We found that EE did not affect the proportion of PV^+^ neurons at P35, nor at P56 (Fig. 5b; P35 naïve n=5 mice, P35 EE n=4 mice, t_(7)_=0.079 p=0.939; P56 naïve n=4 mice, P56 EE n=4 mice, t_(6)_=0.4606, p=0.661, two-tailed unpaired t-test). However, there was a significant experience-dependent increase in the intensity of immunofluorescence signals for PV. Specifically, EE enhanced PV immunofluorescence in PV^+^ neurons lacking a PNN both at P35 and P56 (Fig. 5c; P35 naïve n=503 neurons from 5 mice, P35 EE n=289 neurons from 4 mice; D=1.84, p=7.99×10^-6^; P56 naïve n=316 neurons from 4 mice, P56 EE n=283 neurons from 4 mice, D=0.127, p=0.016, K-S test), pointing to long lasting experience-dependent changes in PV expression in this group of PV^+^ neurons. However, PV^+^ neurons associated with a PNN showed enhanced PV immunofluorescence only at P35 (Fig. 5d; P35 naïve n=382 cells from 5 mice, P35 EE n=378 cells from 4 mice; D=0.173, p=2.467×10^-5^; P56 naïve n=343 cells from 4 mice, P56 EE n=385 cells from 4 mice; D=0.050, p=0.745, K-S test), suggesting that once PNNs accumulate, PV intensity stabilizes.

EE also affected the association of PV^+^ neurons with PNNs. At P35, EE led to an increase in the proportion of PV^+^ neurons with a PNN (Fig. 5c, d top, insets; P35 naïve n=5, P35 EE n= 4 mice; t_(7)_=2.479, p=0.042, two-tailed unpaired t-test). This effect was transient, as it was not observed in EE mice tested at P56 (Fig. 5c, d bottom, insets; P56 naïve n=4 mice, P56 EE n= 4 mice; t_(6)_=0.8727, p=0.416, two-tailed unpaired t-test), suggesting that EE accelerated the transition from lacking a PNN to having a PNN. These results suggest that the expression of PV in PV^+^ neurons lacking a PNN continues to be modulated by experience at least until P56. Differently, the expression of PV in PV^+^ neurons with a PNN reached stable levels by P56 and was not modulated by experience.

In addition to modulating PV immunofluorescence, EE enhanced PNN aggregation, detected as augmented fluorescence intensity of PNNs around PV^+^ neurons. This effect was observed at both P35 and P56, compared to age-matched naïve mice (Fig. 5e; top, P35 naïve n=5, P35 EE n=4 mice; D=0.219, p=2.555×10^-8^; P56 naïve n=4; bottom, P56 EE n=4 mice, D=0.146, p=8.869×10^-4^, K-S test), even in the absence of changes in the proportion of PNNs in GC (Fig. 5e insets; P35 naïve n=5, P35 EE n=4 mice; t_(7)_=0.9735, p=0.363; P56 naïve n=4, P56 EE n=4 mice; t_(6)_=0.7233, p=0.497, two-tailed unpaired t-test). These findings indicate that taste experience at the time of weaning regulates the state of maturation and the capacity for plasticity of PV^+^ neurons. The effect of EE on PV immunofluorescence was long lasting in PV^+^ neurons lacking a PNN, while it was transient in PV^+^ neurons with a PNN, suggesting that once this group of inhibitory neurons accumulates a PNN, the expression of PV becomes insensitive to taste experience. In addition, the effect of EE on PNN aggregation, measured as WFA intensity, was long lasting, indicating that taste experience regulates inhibitory plasticity in GC through distinct mechanisms.

### PNN accumulation regulates the duration of plasticity for sucrose preference

Studies in drosophila and mice show that some forms of taste-dependent plasticity, such as conditioned taste aversion learning and some forms of taste discrimination, persist throughout life (Schier et al., 2019). Here we asked whether the experience-dependent modulation of sweet preference we report is specific to the early postnatal period or whether it persists throughout life. To investigate this, we tested whether our taste exposure paradigm would induce a shift in the sucrose preference curve if started in adult mice. For these experiments, mice were reared on a regular diet until P56 and then exposed to the battery of taste solutions starting at P57 (late exposure; LE; Fig. 6a). Sucrose curves were obtained with BAT four weeks after the end of LE (at P91). Sucrose preference was unaffected by LE (Fig. 6a; naïve n=10, LE n=8 mice; F_(3,173)_=1.130, p=0.338, F test), indicating that the experience-dependent modulation of sucrose preference is developmentally restricted.

**Figure 6.**
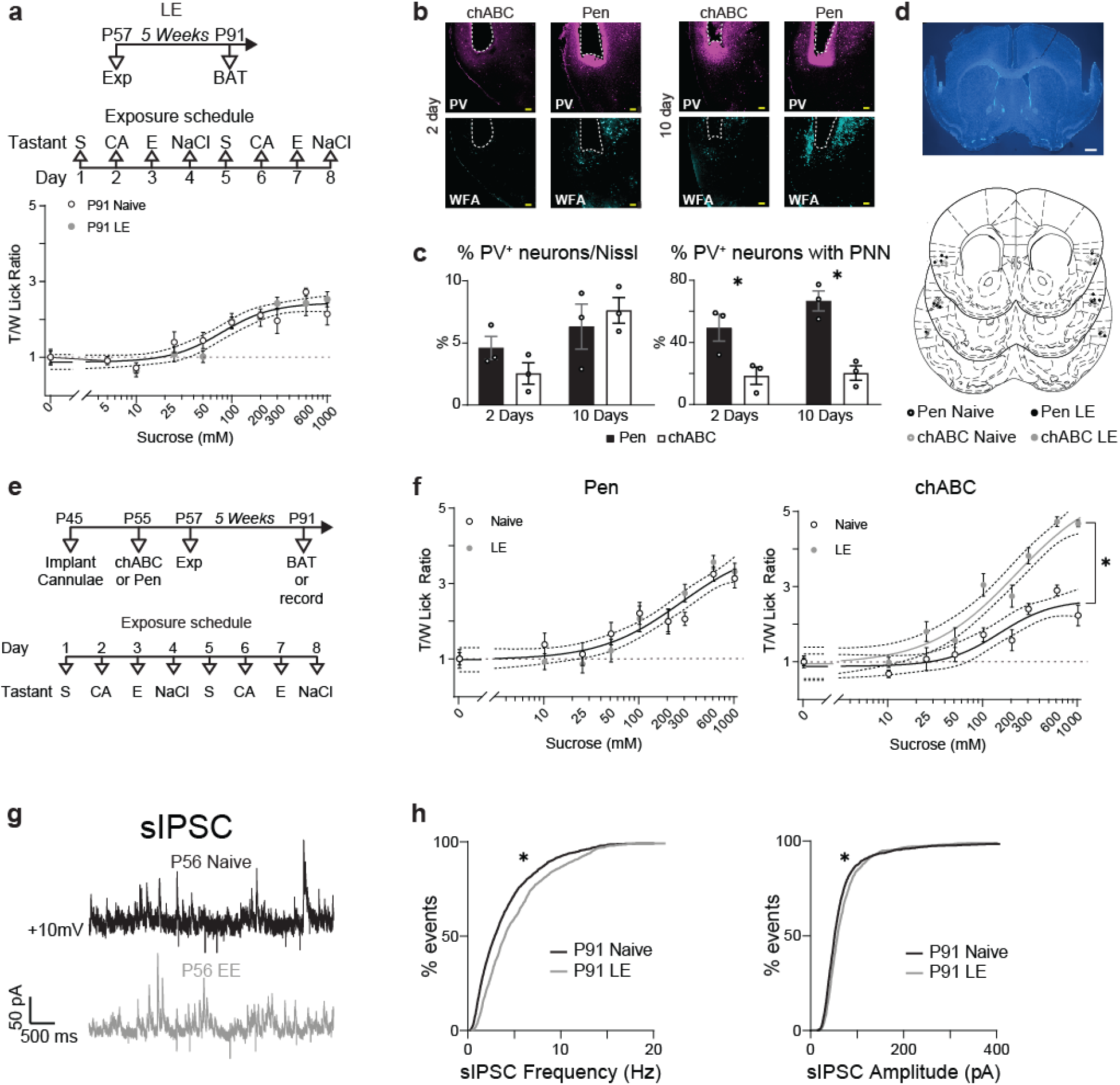
PNN degradation in GC restores sensitivity to taste experience and plasticity of inhibition in adult mice. **a.** Top. Schedule of LE and sequence of tastants. Bottom. Sucrose preference curves for naïve and LE mice tested at P91. A single curve fit both data sets (p=0.338. F test). **b.** Representative images of sections immunostained for PV (magenta) and WFA (cyan) following GC infusion of chABC or Pen. Scale bar=100μm. **c.** Quantification of PV^+^ neurons and PV^+^ neurons associated with a PNN 2- and 10-days post-infusion. Infusion of chABC did not affect the number of PV^+^ neurons (2 days: p=0.178, t-test; 10 days: p=0.566, t-test), however it reduced the number of PV^+^ neurons with a PNN (2 days: p=0.041, t-test; 10 days: p=0.004, t-test). **d.** Top. Representative cannulae placement in GC; brain section stained with Hoechst 33342, scale bar=500μm. Bottom. Cannulae placements for all mice registered to sections of the mouse brain atlas (from Bregma: +1.34mm, +1.10mm, +0.74mm). **e.** Top: experimental schedule for cannulae implant, LE, and BAT. Bottom: sequence of taste solutions. **f.** Left. Sucrose preference curves from P91 mice infused with Pen in naïve (black open circles) or LE (grey filled circles) groups (single curve fit both data sets: p=0.134, F test). Right. Sucrose curves for mice infused with chABC in naïve (black open circles) vs. LE (grey filled circles) group (separate curves fit the datasets: p=2.45×10^-9^, F test). **g.** Sample traces of sIPSCs in mice from naïve (black) and LE (grey) groups, all treated with chABC at P55. **h.** Cumulative distribution of sIPSCs. Left. sIPSC frequency was enhanced chABC-LE mice compared to chABC-naïve (p=3.199×10^-20^, K-S test). Right. sIPSC amplitude also showed a small but significant increase in animals from the chABC-LE group compared to chABC naïve (p=4.407×10^-6^, K-S test). Asterisks indicate p ≤ 0.05.

To determine whether the window for experience-dependent plasticity depends on GC inhibitory circuits and/or on accumulation of PNNs, we used a GC-restricted enzymatic removal of PNNs in adult mice, and asked whether this manipulation could restore the experience-dependent shift in sucrose preference curve. The control group received bilateral intra-GC infusions of pennicillinase (Pen), an enzyme that does not affect PNNs (Pizzorusso et al., 2006; Favuzzi et al., 2017), while the experimental group received intra-GC bilateral infusions of chondroitinaseABC (chABC), an enzyme that breaks the bonds between PNNs proteoglicans, via chronically implanted cannulae. We first quantified the degree of degradation, and the timing of PNN recovery following a chABC injection in GC. A single infusion of chABC in GC at P55 did not affect the proportion of PV^+^ neurons quantified either 2 or 10 days after infusion (Fig. 6b, c left; 2 days: Pen n=3, chABC n=3 mice; t_(4)_=1.631, p=0.178; 10 days: Pen n=3, chABC n=3 mice; t_(4)_=0.625, p=0.566, two-tailed unpaired t-test). However, it was sufficient to degrade PNNs, which did not recover for over 10 days, in line with previous studies (Lin et al., 2008) (Fig. 6b, C right; % PV^+^ neurons associated with a PNN: 2 days: Pen n=3, chABC n=3 mice; t_(4)_=2.986, p=0.041, 10 days: Pen n=3, chABC n=3 mice; t_(4)_=5.794, p=0.004, two-tailed unpaired t-test). Following this validation, separate groups of mice received bilateral GC infusions of Pen or chABC at P55 and were then assigned to either naïve or LE group at P57 (Fig. 6d, e). At the end of the exposure, LE mice returned to their regular diet of chow and water. Sucrose preference curves were obtained at P91 from 4 groups of mice: Pen-naïve, Pen-LE, chABC-naïve and chABC-LE (Fig. 6f). Pen-injected mice showed no change in their sucrose preference curves following LE (Fig. 6f; Pen-naïve n=8, Pen-LE n=8 mice; F_(3,137)_=1.893, p=0.134, F test). However, LE was effective at shifting the sucrose preference curve following in chABC injected mice (Fig. 6f; chABC-naïve n=7, chABC-LE n=9 mice; F_(3,137)_=42.89, p=2.45×10^-9^, F test). These results demonstrate that removal of PNNs locally in GC is sufficient to restore the experience-dependent modulation of sucrose curves in adult mice, indicating that GC is a locus of experience-dependent plasticity, and that mature inhibition in GC contrains experience-dependent changes in sucrose preference.

We next asked whether removal of PNNs in GC also restored the experience-dependent increase of inhibitory synaptic transmission onto PYR neurons. For these experiments, mice received a bilateral injection of chABC at P56, and were then assigned to either the naïve or LE group, beginning at P57. At P91, four weeks after the end of LE, acute slices containing GC were obtained and patch clamp recordings were used to record sIPSCs (Fig. 6e). We observed an increase in sIPSC frequency (Fig. 6g, h; chABC-naïve n=20 cells from 3 mice, chABC-LE n=14 cells from 5 mice; D=0.166, p=3.199×10^-20^, K-S test) and a small, but significant, increase in sIPSC amplitude (Fig. 6g, h; chABC-naïve n=20 cells from 3 mice, chABC-LE n=14 cells from 5 mice; D=0.089, p=4.407×10^-6^, K-S test) in neurons from chABC-LE mice, indicating that removal of PNNs in adult mice allowed for a LE-induced increase in inhibitory synaptic transmission comparable to what we observed following EE.

These data support the interpretation that accumulation of PNNs around PV^+^ neurons in GC limits the capacity for plasticity of inhibitory synapses and prevents the experience-dependent shift in sucrose preference from occurring in adulthood. Thus, the maturation of inhibitory circuits provides a limiting factor that restricts experience-dependent modulation of sweet preference to the first few weeks of postnatal development.

## Discussion

We have shown that taste experience at the time of weaning not only influences the preference for sweet in adults, but also profoundly modulates the postnatal development of gustatory cortical circuits. A convergence of chemosensory signals, nutrient variety and postingestive signals contributes to the experience-dependent shift in sweet preference, highlighting the central role for GC in the integration of sensory information related to feeding. Analysis of licking microstructure supports the interpretation that early taste experience increases the palatability and avidity for sucrose, as EE mice increase the duration of lick bouts and accumulate licks to sucrose faster than their naïve littermates.

We demonstrate *in vivo* that the change in sucrose preference is accompanied by increased spontaneous activity of PV^+^ neuron, reduced spontaneous and sucrose evoked activity in PYR and changes in responsiveness to sucrose concentrations of PYR. Further, our *ex vivo* data indicate that increased inhibitory synaptic drive onto GC PYR neurons is the likely mechanism regulating the responsiveness of these neurons to sucrose. While we observed experience-dependent changes in both excitatory and inhibitory drive at P35, imaging experiments were performed at P56, when excitatory drive had stabilized and only changes in synaptic inhibition were observed. Thus, the long-term effect of EE likely depends on inhibitory synaptic plasticity.

The experience-dependent changes we report are specific for early experience with tastants and nutrients. The exposure paradigm applied to adult mice failed to shift the sucrose preference curve, indicating the presence of a sensitive window during which experience with tastants and nutrients can modulate the preference for sweet. While critical period plasticity has been reported in a different and non-overlapping portion of the insular cortex representing integration of auditory and visual inputs (Gogolla et al., 2014), the prevailing view to date has been that there is no sensitive window for taste (Hensch, 2004). This assumption was based on reports of experience-dependent plasticity throughout life in the gustatory insular cortex (GC) in rodents (Escobar and Bermudez-Rattoni, 2000; Yiannakas and Rosenblum, 2017; Lavi et al., 2018; Haley et al., 2020) and in flies taste neurons (Vaziri et al., 2020). Our data show that taste behaviors can be modulated within restricted developmental windows, provide possible behavioral and neural mechanisms underlying long-term effects of early-life taste experiences in humans (Mennella et al., 2001; Tang et al., 2018; Noble et al., 2019; Volcko et al., 2020), and center GC as a locus of experience-dependent plasticity.

In this study, we focused on GC for its role in taste processing and taste related behaviors. However, taste experience influences the development of other circuits along the taste axis, including the nucleus of the solitary tract (NTS) (Hill et al., 1983). The inputs via cranial nerves terminating in the NTS undergo maturation until P35, and the process remains incomplete in a salt-deficient diet indicating the requirement for early taste experience in the development of the taste system (Sollars et al., 2006; Sun et al., 2017). These results may raise the possibility that the effects we observed rely of indirect GC modulation by other brain regions along the taste system. Indeed, previous work reported that removal of GC does not affect the palatability of tastants, and only mildly impairs simple taste discriminations (Grill and Norgren, 1978; Touzani and Sclafani, 2007; Bales et al., 2015; Blonde et al., 2015; Bales and Spector, 2020). These results were interpreted as evidence that GC is not required for taste identification or palatability. Contrary to these assumptions, our data demonstrate that GC, particularly its inhibitory circuits, play a crucial role in modulating experience-dependent modulation of sucrose preference. Local manipulation of PNNs in GC of adult mice was sufficient to restore the experience-dependent shift in sucrose preference curve, and the increase in synaptic inhibition we observed in younger mice. These results identify accumulation of PNNs around PV^+^ neurons as a mechanism restricting experience-dependent inhibitory plasticity, generalizing findings from other sensory cortical areas to GC (Pizzorusso et al., 2006). In addition, these findings establish GC as an essential circuit driving the establishment of sweet preference during postnatal development.

In summary, our findings highlight the importance of early experience with food and tastants not only for the development of food preferences in adulthood, but also for the postnatal maturation of cortical circuits and sensory processing. They also provide support for the notion that absence of taste experiences early in life, as is the case for tube-fed preterm infants, may have long-term deleterious effects on brain development (Beker et al., 2021).

## Experimental Procedures

### Animals

All experimental procedures followed the guidelines of the National Institute of Health and were approved by the Institutional Animal Care and Use Committee at Stony Brook University. Mice of both sexes were housed in a vivarium on a 12-hour light-dark cycle. All mice were group-housed except for a subset of mice tested in control assessments of baseline tastant consumption (Supplementary Fig. 1a, b; naïve n=4; EE n=3), and for mice used for two-photon imaging experiments (Fig. 3; naïve n=5; EE n=5) which were single-housed following surgery for the prism implant. Experiments were performed during the light cycle. Wild-type C57Bl/6 mice were purchased from Charles River to arrive either as adults or as litters including the lactating mother and pups at postnatal day 5 (P5). PV-cre (*23*) (JAX stock #017320) and Ai14 (*24*) (RRID: IMSR_JAX:007908) were ordered from Jackson Laboratory. The PV-cre;Ai14 mice were heterozygous for both Cre and Lox-Stop-Lox-tdTomato alleles and were bred in house by crossing female homozygous PV-cre mice with males homozygous for the Ai14 reporter gene.

### Behavioral procedures

#### Exposure

Exposure began at P22 for the Early Exposure (EE) group and P57 for the Late Exposure (LE) group. The animal’s homecage water bottle was replaced with a tastant solution following an 8-day schedule (Fig. 1a; 150mM sucrose (S) on days 1 and 5, 20mM citric acid (CA) on days 2 and 6, Ensure (E) on days 3 and 7, and 100mM salt (NaCl) on days 4 and 8). Ensure was delivered in a bowl to eliminate the possibility of clogging the spout. Mice had ad lib access to chow. Control mice (naïve) remained on a diet of ad lib chow and water. For the calorie-free EE, the homecage water bottle was replaced with a tastant solution in the following 8-day schedule: 8mM saccharin on days 1 and 5, 20mM citric acid (CA) on days 2 and 6, 100mM monosodium glutamate (MSG) on days 3 and 7, and 100mM salt (NaCl) on days 4 and 8. For experiments in which animals were given 8 days of either sucrose, salt, or Ensure, they were treated as above, but with the appropriate taste solution.

#### Tube feeding

Tube feeding began on P22 and ended on P29. Polyurethane tubing (0.63×1.02mm) was marked with the length from the mouth to the animal’s sternum and attached to a 10mL Hamilton syringe. Infusates were either water or Ensure. The concentration of Ensure was determined by calculating the amount of Ensure consumed during 1-2 hours access in the anosmia experiment (see below) to approximate the calorie intake by mice of a certain body weight. The Ensure was then mixed to be more concentrated in order to accommodate the smaller volume given by tube-feeding. The concentrated Ensure was administered up to three times/day. The exact concentration was calculated for each animal’s body weight on each day. The highest concentration was 6g/mL. Animals were scruffed and the tubing gently inserted into the mouth and down the esophagus until the necessary length of tubing was inserted. Infusate was then slowly administered by hand at approximately 0.1mL/10g bodyweight. Animals were monitored after infusions for signs of aspiration or discomfort, neither of which were observed.

#### Intranasal instillation for anosmia

Intranasal instillations of ZnSO_4_ or saline were performed as previously described with modifications for weanlings (Czarnecki et al., 2011). To determine the appropriate length for the infusion apparatus and volume of infusate, a pilot experiment was conducted in which AlexaFluor488 dye transport was observed from the nasal epithelium to the olfactory bulbs without expulsion or aspiration of infusate. For experimental animals, P21 mice were lightly anesthetized with a cocktail of ketamine (70mg/kg) and dexmedetomidine (1mg/kg). Eppendorf microloaders were attached to a 1μL Hamilton syringe. Animals were placed on their side while a microloader was threaded 5mm into the naris. Animals were then rotated onto their back while 4μl infusate (ZnSO_4_ [155mM] or saline) was slowly administered. The mice remained in this position for 20 minutes, then they were rotated to their opposite side. After a 5-minute delay, the same procedure was conducted for infusion into the opposite naris. Animals were then given 1mg/kg antisedan to reverse the dexmedetomidine.

For delivery of Ensure for 8 days in ZnSO_4_ or saline treated mice, individual mice were transferred from their homecage to a holding cage containing standard bedding for 1-2 hours/day with access to a small bowl of Ensure. The amount of Ensure consumed on each day was measured by weighing the bowl before and after consumption. This procedure allowed us to measure the amount of Ensure consumed by individual mice while maintaining group housing.

#### Brief Access Test (BAT)

Procedures were adapted from Glendinning et al (Glendinning et al., 2002). Sucrose preference curves were collected using a commercially available gustometer (Med Associates Inc, Davis Rig for Mouse-16 Bottle). The apparatus consisted of a chamber (14.5 cm wide, 30 cm deep, 15 cm tall), a motorized moving table to deliver multiple taste stimuli, a motorized shutter door to allow temporary delivery of tastants, and a dedicated computer to control tastant delivery and to record the timing of licks. Licks were detected by a high frequency AC contact circuit activated by tongue contact with the sipper tube. The apparatus was cleaned with 70% ethanol between mice. The number of licks, the latency to the first lick, and inter-lick intervals (ILIs) were recorded for each trial. For analysis, ILI <60ms were removed to exclude double contacts occurring from a single tongue protrusion (Glendinning et al., 2002). Water restriction began on day 1 of training. Water restriction parameters were mild to minimize thirst-driven consumption and to maximize licking due to the appetitive nature of sucrose. Mice were weighed daily to ensure maintenance of at least 80% of their initial body weight. The table below summarizes the schedule of training and access to water.

| **Day** | **Training (all sessions 30min)** | **Access to Water** |
| --- | --- | --- |
| 1 | Context Habituation | Water bottle removed after training session |
| 2 | Spout Training | Only during training |
| 3 | Shutter Training | 10min at the end of the day |
| 4 | Collection First Sucrose Curve | 5min water 2h prior to testing & 10min at the end of sucrose curve collection |
| 5 | Collection Second Sucrose Curve | 5min water 2h prior to testing; returned to ad libitum food and water at the end of sucrose curve collection |

On day 1 of training, mice were habituated to the Davis rig gustometer in a 30min session where the shutter door remained opened and granted access to a sipper tube connected to a bottle containing water. At the end of the first session, the water bottle was removed from the homecage and water restriction began. On day 2, mice were returned to the gustometer for a 30min spout training session, which began on the first lick. Once the mouse was placed in the chamber, the shutter opened, granting access to a sipper tube connected to a bottle containing water. Mice were allowed to lick ad libitum during the 30min session. On day 3, mice were familiarized with the trial structure and the opening and closing of the shutter door (shutter training). These 30min sessions consisted of trials in which the shutter door opened, granting access to a sipper tube connected to a bottle containing water. If the mouse did not lick, the shutter door closed after 15s, followed by a 5s inter-trial interval (ITI). If the mouse did lick, the shutter door remained open for 10s following the first lick. At the end of the 10s interval, the shutter door closed. During the 5s ITI the motorized table moved in order to present a different bottle containing water. At the end of day 3, mice had 10min access to a water bottle in their homecage. On days 4 and 5, mice were weighed and then given 5min access to a water bottle in the homecage 2h prior to being placed in the gustometer for the acquisition of sucrose preference curves. At the end of day 4, mice had 10min access to a water bottle in the homecage, while at the end of day 5 the experiment was concluded, and mice were returned to ad libitum access to water. On test days (4 and 5), the trial length was 10s with a 5s ITI. The shutter door closed after 10s if the mouse did not lick.

To obtain sucrose preference curves, 6 bottles containing different concentrations of sucrose were presented in a pseudorandom order. The range of concentrations were presented as a block and the software randomized the presentation (without replacement) within each block. The sucrose concentrations presented on sucrose curve 1 were (in mM): 0, 5, 10, 50, 100, 600, and those presented on sucrose curve 2 were (in mM): 0, 5, 25, 200, 300, 1000. Sucrose curve 1 or 2 was presented on either day 4 or day 5 of testing, ensuring that presentation was counterbalanced within groups. Mice in the EE group were P56 on day 4; mice in the LE group were P91 on day 4. For BAT for assessing salt preference, the salt concentrations presented on salt curve 1 were (in mM): 0, 10, 50, 100, 250, 500, and those presented on salt curve 2 were (in mM): 0, 25, 75, 400, 750, 1000. Salt curve 1 or 2 was presented on either day 4 or day 5 of testing, ensuring that presentation was counterbalanced within groups. For BAT assessing saccharin preference, the saccharin concentrations presented on saccharin curve 1 were (in mM): 0, 0.2, 2, 10, 20, and those presented on saccharin curve 2 were (in mM): 0, 0.6, 6, 20. Saccharin curve 1 or 2 was presented on either day 4 or day 5 of testing, ensuring that presentation was counterbalanced within groups. For BAT following chABC or Pen infusion presented in Fig. 6F, 5mM sucrose was omitted from testing. Mice had to participate in at least 20 trials to be included in the final analyses. For the sucrose preference curves collected at P56, 4 out of a total of 93 mice used were removed from analysis because their average lick rate to water was less than 1Hz.

#### Behavioral protocol for 2P imaging

All procedures were conducted on a custom designed behavioral apparatus controlled with codes written in LabView (National Instruments). Events were recorded on an Intan RHD USB Interface Board (RHD2000, Intan Technologies). Licks were detected using an electronic circuit (Slotnick, 2009). Following recovery from prism implantation surgery (see below), mice were placed on water restriction for one week (85% body weight, 1-1.5mL/water daily). Next, they were habituated to perform 5 dry licks to receive one drop of water (4μl) from a motorized spout (X-LSM, Zaber Technologies). The intertrial interval for each session was 10 ± 4s. If an animal failed to perform 5 dry licks within 3s, the spout retracted and an additional timeout of 10s was triggered. After 4 days of habituation, two photon imaging with sucrose concentrations began. Mice needed to perform 5 dry licks to receive one drop (4μl) of either sucrose or water. The following concentrations of sucrose were used (in mM): 0 (water), 25, 50, 100, 200, 600. On each trial, a water rinse (4μl) was given 8.3 ± 0.25s after each taste. Tastants were delivered using gravity fed solenoid valves (Lee Company) controlled by a ValveLink8.2 controller (Automate Scientific). Equal numbers of trials for each taste solution were delivered in a pseudorandom order. Mice were allowed to continue performing the task until they stopped licking (total trials, naïve: 109.9 ± 5.1, EE: 96.1 ± 6.1; t-test p>0.05, t=1.673, df=25). The average number of licks across all tastants was comparable across conditions (naïve: 11.7+/- 0.1; EE: 11.9 +/- 0.1; t-test p>0.05, t=0.7406, df=25), indicating comparable volume consumption across conditions.

#### Buried Food Test

Mice were treated with intranasal instillations at P21. Behavior tested procedures were modified from (Yang and Crawley, 2009) and were video recorded. Pre-exposure to Froot Loops was required and began on P22 in their homecage for 3 days (P22-P24) before testing. Testing took place at P25 or P28. The day before testing, chow was removed from the homecage at approximately 5pm for an overnight food restriction. On test day, mice were individually transferred to a clean shoebox cage containing at least 3cm of bedding and allowed to explore for 5min. They were then removed from the test cage and returned to their homecage while one-quarter to one-third piece of Froot Loop was buried in the test cage. They were then returned to the test cage and placed in the half of the cage opposite to where the target Froot Loop was buried. The time to sniff in the correct location, dig in the correct location, and retrieve the pellet were recorded. Animals were allowed to explore and attempt to find the pellet for 15min (900s), at which point timeout occurred, the pellet was unburied, and they were allowed to consume it. All mice quickly found and ate the pellet once unburied. Analysis was done on the time to dig in the location of the pellet.

#### Habituation/Dishabituation

Anosmia and recovery from anosmia were assessed with a non-associative behavioral task designed to assess the mouse’s investigation of a novel odor (modified from (Kass et al., 2013). Mice were individually transferred from their homecage to a clean empty cage of the same size. On habituation trials, a hexagonal weigh boat containing a small piece of filter paper dipped in mineral oil was presented on the wire cage top such that the mouse could sniff and explore but not touch. Three 60s habituation trials with mineral oil were presented with a 2min ITI. In the final trial (also 60s), mice were presented with a hexagonal weigh boat containing a small piece of filter paper dipped in methyl valerate (1:500 concentration diluted in mineral oil) or isoamyl acetate (1:500 concentration diluted in mineral oil). Time spent sniffing was included if the mouse was standing in the third of the cage containing the weigh boat and engaged in active sniffing directed towards the weigh boat.

### Surgery

#### Implants of infusion cannulae

Mice aged P45-48 were anesthetized with an intraperitoneal (IP) injection of a cocktail containing 100mg/kg ketamine and 10mg/kg xylazine. Bupivacaine (2.5mg/ml, ~0.1mL) was administered under the scalp for local anesthesia. Craniotomies were made over the left and right GC at the following coordinates: +1.0mm anterior from Bregma, 3.25mm lateral from midline. Stainless steel guide cannulae (26 gauge, 5.0mm, Plastics One) were lowered 1.8mm vertical from the cortical surface, fixed to the skull with acrylic dental cement (Stoelting). A metal head post was implanted posterior to the cannulae for head restraint during infusion. Once the dental cement was completely dry, dummy cannulae were inserted into each guide cannula, and animals were allowed to recover for at least 1 week before the infusion. For the experiments examining the time course of recovery from the chABC (Fig. 6), surgical procedures were unilateral with a single cannula implant in the left GC. For experiments shown in Fig. 6d-f cannula implants were bilateral.

#### Injection of viral construct and prism implant

In preparation for two photon imaging, PV-cre;Ai14 mice underwent two surgeries. The first was performed at P20 or P21. Mice were anesthetized with the ketamine/xylazine cocktail described above and a craniotomy was made over the left GC at the following coordinates: +0.8mm from bregma, ~3.5mm lateral from midline (close to the lateral suture). A pulled glass pipette attached to a 10μl Hamilton syringe was backfilled with a solution containing a viral construct carrying gCaMP7f (AAV1-syn-jGCaMP7f-WPRE, titer 3×10^13^ gc/mL Addgene cat no 104488-AAV1). To reach the appropriate titer, the stock virus solution was diluted 1:4 in sterile saline. Injections were delivered at 1.75mm and 1.9mm below the pial surface. The syringe tip was slowly lowered in the brain to the desired depth and a total volume of 225nl was infused at a rate of 1nl/sec, using a microinjection syringe pump (UMP3T-1, World Precision Instruments). The pipette tip was left in place for 10min after each infusion before being slowly retracted. The craniotomy was filled with silicone (Dow Corning 200) and the skin sutured closed. Mice were given an injection of lactated Ringer’s solution, placed back into the homecage and allowed to recover overnight on a heating pad. The exposure paradigm began 1-2 days later at P22. A second surgery to implant prisms was performed at P42-P44. Mice were anesthetized as described above and administered subcutaneous injections of dexamethasone (2mg/kg) and carprofen (5mg/kg) before beginning the surgery. Bupivacaine (2.5mg/ml, ~0.1mL) was administered under the scalp for local anesthesia. The skin above the skull and overlying the left temporalis muscle was removed and the skull cleaned. Portions of the temporalis muscle were removed and a ~2.2 x 2.2mm craniotomy was opened on the lateral surface of the skull with a dental drill using the middle cerebral artery as a landmark for GC. A durotomy was performed using fine forceps. The prism assembly was lowered into place and secured with Vetbond and black dental cement. A custom headpost was affixed to the skull using black dental cement. Prisms were fabricated by cutting 3 x 3mm coverslips (Warner Instruments, cat no 64-0728) to approximately 3 x 2mm size using a diamond tipped etcher. Coverslips were cleaned with ethanol, dried, and glued onto the brain facing surface and aluminum coated hypotenuse of a 2mm microprism (Tower Optical, cat no MPCH-2.0) using UV curable glue (Norland, cat no NOA-61). Mice were given carprofen (5mg/kg) daily for 5 days after the surgery and allowed to recover for at least 1 week before beginning water restriction and behavioral and imaging studies.

#### Injection of chABC for slice electrophysiology

Wild-type mice at age P54-56 were anesthetized with the ketamine/xylazine cocktail described above. Bupivacaine (2.5mg/ml, ~0.1mL) was administered under the scalp for local anesthesia. Craniotomies were made over the left and right GC at the following coordinates: +1.0mm anterior from Bregma, 3.25mm lateral from midline. A pulled glass pipette was attached to a nanoject pressure injector (Drummond nanjoect II) and filled with chABC in its vehicle (described below). The lipophilic tracer, DiI, was applied to the outside of the glass pipette in order to later determine the location of chABC injection. The pipette was lowered 2.2mm below the pia. Approximately 0.25μl total volume was injected per side over 10 minutes. The pipette tip was left in place at least 10min after the end of the injection. The skin was then sutured closed, and the animal recovered on a heating pad. Mice were then returned to their homecage and the LE protocol began 2 days later. Only animals with DiI fluorescence in GC were included in the final analysis.

### Enzyme infusions

P55 mice were head-restrained, the dummy cannulae were removed, and an injection cannula (33 gauge, 5.0mm with 0.5mm projection, Plastics One) was inserted into each guide cannula. Injection cannulae were attached to 10μl Hamilton syringes via polyethylene tubing from A-M systems (Sequim, Wa). The injection cannulae extended an additional 0.50mm from the tip of the guide cannulae in order to reach GC. Bilateral infusions of 100U/mL chABC (or 100U/mL Pen) were delivered at a flow rate of 0.1μl/min for a total volume of 0.25μl per side. Infusion rate was controlled by a pump (Harvard Apparatus). To preserve enzymatic activity, chABC and Pen were dissolved in a vehicle solution containing 50mM Tris-HCL, 60mM sodium acetate, and 0.2% BSA (Pizzorusso et al., 2006; Favuzzi et al., 2017). Following infusion, the injection cannulae were left in place for 2min to allow the solution to diffuse from the cannula tips. The dummy cannulae were subsequently reinserted into the guide cannulae. The late exposure paradigm began 2 days later, on P57.

### Two-photon imaging

Imaging experiments began after 4 days of habituation to the behavioral apparatus. Images were acquired on a two-channel movable objective microscope (MOM, Sutter) using a resonant scanner controlled by MScan (Sutter) mounted with a 10x super apochromatic objective (Thorlabs, 0.5NA, 7.77mm WD). Fluorophores were excited using a Ti:Sapphire femtosecond laser (Coherent) tuned to 980nm with a laser power of 100-250mW at the front of the objective, and emission was collected with using photomultiplier tubes (Hamamatsu). Recording was triggered in episodes ~4s before the start of each trial. For each episode, 520 frames were acquired at 31Hz in both the red and green channels. After each session a 40μm z stack centered on the imaging plane was obtained for both channels to aid in identification of PV^+^ neurons. All mice with sufficient optical window clarity for imaging were trained and included in the analysis.

### Electrophysiology

P34-36, P55-57, or P90-92 mice were anesthetized with isoflurane using the bell jar method and rapidly decapitated. The brain was dissected in ice cold, oxygenated standard artificial cerebrospinal fluid (ACSF) containing, in mM: 126 NaCl, 3 KCl, 25 NaHCO3, 1 NaHPO4, 2 MgSO_4_, 2 CaCl2, 14 dextrose with an osmolarity of 313-317mOsm with a pH = 7.4 when bubbled with carbogen (95% oxygen, 5% carbon dioxide). Coronal slices (350μm) containing GC were prepared using a fresh tissue vibratome (Leica VT1000), allowed to recover in 34°C ACSF for 20min, and then brought to room temperature for 30min before beginning experiments. Individual slices were transferred to the recording chamber and perfused with an ACSF solution optimized to increase spontaneous activity (*38*), in mM: 124 NaCl, 3.5 KCl, 26 NaHCO3, 1.25 NaHPO4, 0.5 MgCl2, 1 CaCl2, 14 dextrose, maintained at 34°C with an inline heater (Warner). Whole-cell patch clamp recordings were obtained from visually identified pyramidal neurons under DIC optics using borosilicate glass pipettes with resistance of 3–5MΩ and filled with a cesium sulfate-based internal solution containing, in mM: 20 KCl, 100 Cs-sulfate, 10 K-HEPES, 4 Mg-ATP, 0.3 Na-GTP, 10 Na-phosphocreatine, 0.2% biocytin (V_rev_[Cl^−^1] = −49.3 mV). The pH was adjusted to 7.35 with KOH; osmolarity was adjusted to 295mOsm with sucrose. The sodium channel blocker QX314 (3mM, Tocris Bioscience) was used to stabilize recordings during prolonged depolarization.

Spontaneous postsynaptic currents (sPSCs) were recorded in voltage clamp by holding neurons at three different holding potentials (Tatti et al., 2017). For each neuron, current vs. holding voltage (V_hold_) functions were used to identify the voltage that better isolated the current of interest, which were used for analysis of spontaneous events’ amplitude and frequency (*12*, *38*). To isolate spontaneous inhibitory postsynapic currents (sIPSCs), recordings were obtained around the reversal potential for AMPA and NMDA mediated currents (in mV: +5, +10, +15). Cumulative distribution functions were created from 100 sIPSCs from each recorded cell. Spontaneous excitatory synaptic currents (sEPSCs) were recorded at three different holding potentials around the reversal potential for chloride (in mV: −55, –50, −45). Cumulative distribution functions were created from 35 sEPSCs from each recorded cell.

Recordings of input-output functions and dynamic input resistance (DIR) were performed as previously described (Swanson et al., 2021) in current clamp with a K-Gluconate-based internal solution containing, in mM: 100 K-Gluconate, 20 KCl, 10 K-HEPES, 4 Mg-ATP, 0.3 Na-GTP, 10 Na-phosphocreatine, and 0.4% biocytin, pH was adjusted to 7.35 with KOH; osmolarity was adjusted to 295mOsm with sucrose. Current steps (700ms) of increasing intensity (−100 to 600pA at 50pA steps) were injected into the cell with a 10s inter–sweep interval. DIR is reported as the slope of the current-voltage curve for steps below action potential threshold. Input-output curves represent the average action potential firing frequency for suprathreshold current step for each experimental condition. Neurons with a series resistance >15MΩ or that changed >20% during recording were excluded from the analysis. Neuron identity was confirmed post-hoc with immunohistochemistry aimed at reconstructing neuron morphology, determining location, and assessing expression of the GABA neuron marker GAD67.

### Immunohistochemistry

#### Validation of cannulae placement or gCaMP7f injection sites

Mice were deeply anesthetized and transcardially perfused with phosphate-buffered saline (PBS), followed by perfusion with 4% paraformaldehyde (PFA) in PBS. Brains were dissected out and postfixed in 4% PFA at 4°C for 3-24h followed by cryoprotection in a PBS-buffered sucrose (30%) solution until brains were saturated (~36 h). Thin (50μm) coronal brain sections containing the GC were cut on a vibratome (VT1000, Leica). Brain sections were first washed in PBS (3 × 5min) at room temperature (RT), then incubated with the nuclear stain, Hoechst 33342 (1:5000, Invitrogen, H3570) in PBST (0.1% Triton X-100) for 30min at RT. Sections were then washed with PBS (3 x 10min) and mounted onto glass slides with Fluoromount-G. All sections containing cannulae tracks or gCaMP7f signal were mounted and imaged on a fluorescence microscope (Olympus BX51WI) at 4x magnification.

#### PV^+^ neuron and PNN labeling

Mice were deeply anesthetized and transcardially perfused with PBS, followed by perfusion with 4% PFA in PBS. Brains were dissected out and postfixed in 4% PFA at 4°C for 3-24h followed by cryoprotection in a PBS-buffered sucrose (30%) solution until brains were saturated (~36h). Thin (50μm) coronal sections containing GC were cut on a vibratome (VT1000, Leica). Brain sections were first washed in PBS (3 × 5min) at room temperature (RT) and then blocked in 5% normal goat serum (NGS) and 5% bovine serum albumin (BSA) in PBST (0.5% Triton X-100) for 1h at RT. These preparatory steps were followed by incubation with primary antibodies overnight at 4°C in a solution containing 2% NGS and 1% BSA in PBST (0.1% Triton X-100). The following antibodies were used for quantification and analysis of fluorescence intensity of PV^+^ neurons and PNNs: mouse anti-parvalbumin (1:2000, Swant 235, RRID: AB_10000343), and biotinylated wisteria floribunda lectin (1:500, Vector Laboratories B-1355-2). Sections were then washed with PBS (3 × 10min) and incubated with the following fluorescent secondary antibodies at RT for 2h: Goat anti-mouse Alexa Fluor 488 (1:200, Thermo Fisher A-11029), Streptavidin Alexa Fluor 647 (1:500, Thermo Fisher S32357) and counterstained with neuronal-targeting fluorescent Nissl (Neurotrace530/615, 1:200 Thermo Fisher N21482). After washing with PBS (3 × 10min), sections were mounted onto glass slides with Fluoromount-G (Southern Biotech). Images were obtained with a laser-scanning confocal microscope (Olympus Fluoview) at 10x magnification. Fluorescence signal quantification was performed on two sections of GC spaced at 150μm from each animal. Quantification of the number of PV^+^ neurons and PNNs as well as of fluorescence intensity was obtained in ImageJ using the Pipsqueak AI macro from Rewire Neuro, Inc (Slaker et al., 2016). Individual ROIs for each PV^+^ neuron or PNN were automatically created and verified by an experimenter who was blind to the experimental condition. All single- or double-labeled neurons or PNNs were detected and analyzed. The proportion of PV^+^ neurons associated with PNNs and the intensity of each ROI was also assessed. The proportion of PV^+^ neurons was quantified relative to the neuron-specific Nissl-labeled counterstain (Neurotrace). Quantification of neurotrace-positive cells was performed using the ImageJ plugin image-based tool for counting nuclei (ITCN; Center for Bio-image Informatics, University of California, Santa Barbara).

#### Labeling of recorded neurons

After electrophysiological recordings, slices were post-fixed in 4% PFA for at least 1 week. They were then washed in PBS (3 × 5min) at room temperature (RT) and then were blocked in 5% NGS and 5% BSA in PBST (1.0% Triton X-100) for 2-4h at RT, followed by incubation with primary antibodies overnight at 4°C in a solution containing 1% NGS and 1% BSA in PBST (0.1% Triton X-100). The following antibodies were used: streptavidin Alexa Fluor-568 conjugate (1:2000, Invitrogen, S11226), mouse anti-GAD67 (1:500, MilliporeSigma, MAB5406, monoclonal). Sections were then washed with PBS (3 × 10min) and incubated with the following fluorescent secondary antibodies at RT for 4h: Goat anti-mouse Alexa Fluor 488 (1:200, Thermo Fisher A-11029), and counterstained with a nuclear stain, Hoechst 33342 (1:5000, Invitrogen, H3570). After washing with PBS (3 × 10min), sections were mounted onto glass slides with Fluoromount-G. Sections were imaged with a laser-scanning confocal microscope (Olympus Fluoview) at 10x magnification for validation of location within the GC and at 40x to determine possible colocalization with GAD67.

### Calcium Imaging

Episodes were downsampled from 31Hz to 6.2Hz (ImageJ, grouped Z projection) before downstream processing. Rigid motion correction was performed using NormCorre (Pnevmatikakis and Giovannucci, 2017) and ROIs, calcium traces and deconvolved activity were extracted using the constrained non-negative factorization algorithm (CNMF) (Pnevmatikakis et al., 2016). The automatically detected ROIs were manually curated by inspecting traces and ROI shapes. The deconvolved activity, representing putative spikes, was used in all downstream analysis. PV^+^ neurons were identified using the 40μm z stack obtained at the end of each imaging session. The channels (green and red) were separated, and each stack was collapsed (ImageJ, Z projection). The extracted gCaMP ROIs were overlaid onto the green channel and shifted if necessary to account for potential drift across the motion correction/session. ROIs were then overlaid onto the red (PV) channel and a 4-pixel annulus was drawn around each ROI (ImageJ, make band). The ratio of the pixel intensity within the ROI to that within the band was computed, and ROIs were sorted based on this ratio (ImageJ, measure). Cells in this sorted list were inspected to determine a cutoff ratio for identifying them as PV^+^. All cells above the cutoff value were manually inspected to obtain final PV labels. Cells deemed PV^−^ were grouped as putative PYR. PV^+^ cells consisted of ~5% of imaged cells, consistent with previous studies using cell counting and other approaches (Gonchar et al., 2007; Kim et al., 2017).

### Two photon data analysis

Putative spikes from calcium imaging data were aligned to taste delivery and separated into 1s bins. A neuron was considered taste responsive if the mean activity in a 1s baseline before licking onset was significantly different from activity in one of the three bins after taste delivery (rank sum P<0.05, corrected for 6 tastes/3 bins). Qualitatively similar results were obtained when using a different 1s baseline. Proportions of responsive neurons were compared using a chi-square test. For curve fitting analysis, mean activity for each sucrose concentration was identified for each 1s bin and fit with a linear function. For each fit, 95% confidence intervals (CIs) were calculated, and a bin was determined to have a significant fit if the CI did not include zero. Neurons were considered to have a significant linear response if at least one bin in the 3s sampling window displayed a significant fit. For the baseline, calculations were performed on an equivalent number of bins, and the proportions of significant fits were compared between naïve and EE using a chi-square test. For the neurons with a significant fit, the absolute maximum slopes were compared in both the baseline and sampling windows using an unpaired two-tailed t-test.

### Data analysis and statistics

Analysis of data obtained with the Brief Access Test (BAT) were performed using GraphPad Prism 9, version 9.2.0. A tastant/water lick ratio was calculated by quantifying the number of licks in each trial and dividing that value by the average licking for all water (0mM sucrose) trials. Thus, a tastant/water lick ratio of 1.0 occurred when the licking was similar to licking for water, indicating no difference in preference. Data analysis was done by fitting sigmoid functions to each data set with 3 independent parameters (Top, IC50, HillSlope) and 1 shared parameter (Bottom). Data points and error bars represent mean ± standard error of the mean (SEM). For salt preference curves presented in Supplementary figure 3, the sigmoid functions were fit with all 4 independent parameters (Bottom, Top, IC50, HillSlope) due to the complex nature of salt preference and avoidance. Sigmoid curves followed the equation:

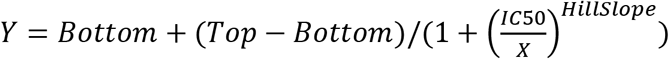

Sigmoid fits were plotted as fitted curve ± 95% confidence bands. Fitted curves were analyzed for statistical differences using the extra sum-of-squares F test (“F test”). For BAT experiments involving naïve mice at P56, the same group of 26 naïve mice were used for 7 different comparisons across multiple experiments (P56 naïve vs P35 and P91 in Fig 1C; vs P56 EE in Fig 1D; vs sucrose only EE, salt only EE, and Ensure only EE in Fig 1E; vs calorie-free EE in Fig 2A). For these experiments, p value for significance was set to 0.0071 to correct for multiple comparisons.

For lick structure analysis, analyses were completed using GraphPad Prism 9, version 9.2.0. Licking bouts were defined as a minimum of 3 licks, each occurring within 250ms of the next. Sessions were excluded if the mouse made fewer than 200 licks. Licking bout data were analyzed with Mann-Whitney U test, and accumulated licks were analyzed with two-sample K-S tests. Licks were sorted into bouts using custom Python scripts. Histograms and the Shapiro-Wilks test were conducted to test normality of the data prior to running post-hoc nonparametric statistical tests to compare data. Accumulated licks were calculated using the following equation:

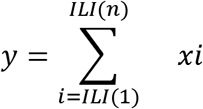

Cumulative distributions of sIPSCs amplitude and frequency were obtained in Igor (WaveMetrics) and Clampfit (Molecular Devices) with the event detection template search. Statistical comparisons were assessed with a two-sample K-S test. To avoid statistical complications of large sample sizes for these data sets (>1000), we applied a more stringent p value of 0.01 for significance. Averages presented represent mean ± SEM. Cumulative distributions of fluorescence intensity were extracted from Pipsqueak plugin for ImageJ and analyzed with a two-sample K-S test; intensity averages were analyzed with two-tailed t-tests.

## Acknowledgments

This work was supported by

National Institutes of Health grant R01DC019827 to AM

National Institutes of Health grant R01DC013770 to AM and AF

National Institutes of Health grant R01DC015234 to AF and AM

National Institutes of Health grant UF1NS115779 to AF and AM

National Institutes of Health Fellowship F32DC018485 to HCS

National Institutes of Health Fellowship F30DC019523 to JFK

## Author Contributions

Designed experiments: HCS, JFK, MI, LAC, AF, AM

Performed experiments: HCS, JFK, MI, LAC

Analyzed data: HCS, JFK, MI, LAC

Supervised experiments and analysis: AF, AM

Writing: HCS, JFK, MI, LAC, AF, AM

## Supplementary Figures

**Supplementary Figure 1:**
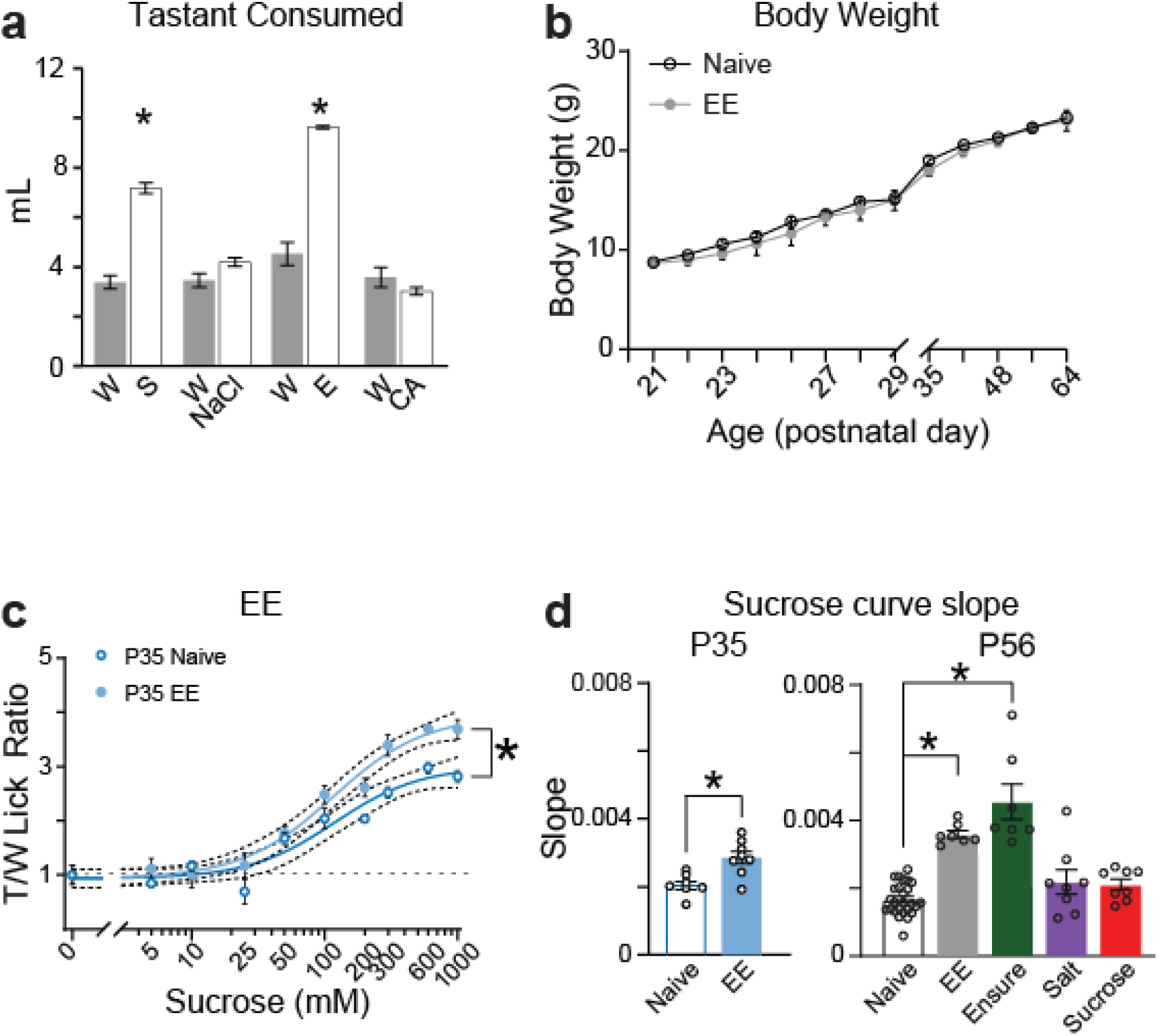
Additional analysis of the effects of EE. **a-b.** To compare baseline consumption of water and of each taste solution, separate groups of mice were single-housed and split into groups given access to either taste solutions (tastant group) or to water (water only group) over subsequent days in their homecage water bottle. **a.** Mice with access to Ensure or sucrose consumed more fluid compared to mice with access to water (water only n=4 mice, tastant-exposed n=3 mice; (sucrose: t_(5)_=11.36, p<0.001), two-tailed t-test; Ensure: t_(5)_=12.96, p<0.001, two-tailed t-test). Mice with access to NaCl and citric acid consumed volumes comparable to mice with access to water (water only n=4 mice, tastant-exposed n=3 mice, NaCl: t_(5)_=2.33, p=0.07, two-tailed t-test; citric acid: t_(5)_=(1.501), p=0.19, two-tailed t-test). Thus, mice consumed all tastant in the exposure paradigm comparably to or more than water. **b.** Comparison of weight gain between naïve mice and those in the EE group showed no differences in body weight either during the exposure paradigm (P21-P30) or during the six weeks following the end of EE (F_(1,5)_=0.3662, p=0.57, 2-way RM ANOVA). **c.** Sucrose curve for P35 naïve (open blue circles) and EE (filled light blue circles) mice. Separate curves were needed to fit the datasets (P35: naïve n=7, EE n=8 mice; F_(3,143)_=14.94, p=1.64×10^-08^, F test). **d.** Plot of the slope of sucrose curves obtained from different mice engaged in the distinct EE paradigms. One-way ANOVA (F_(4,50)_=28.03; p=3.094×10^-12^, 1-way ANOVA) and post hoc Bonferroni multiple comparisons tests revealed no differences in the slope of curves from mice from naïve (n=26 mice); 8 days of NaCl (n=8 mice, p=0.319) or 8 days of sucrose (n=7 mice, p=0.480), while sucrose curves from mice in the EE group (n=7 mice, p=2.618×10^-7^) or those exposed to 8 days of Ensure (n=7 mice, p=3.639×10^-12^) were steeper. Asterisks indicate p ≤ 0.05.

**Supplementary Figure 2.**
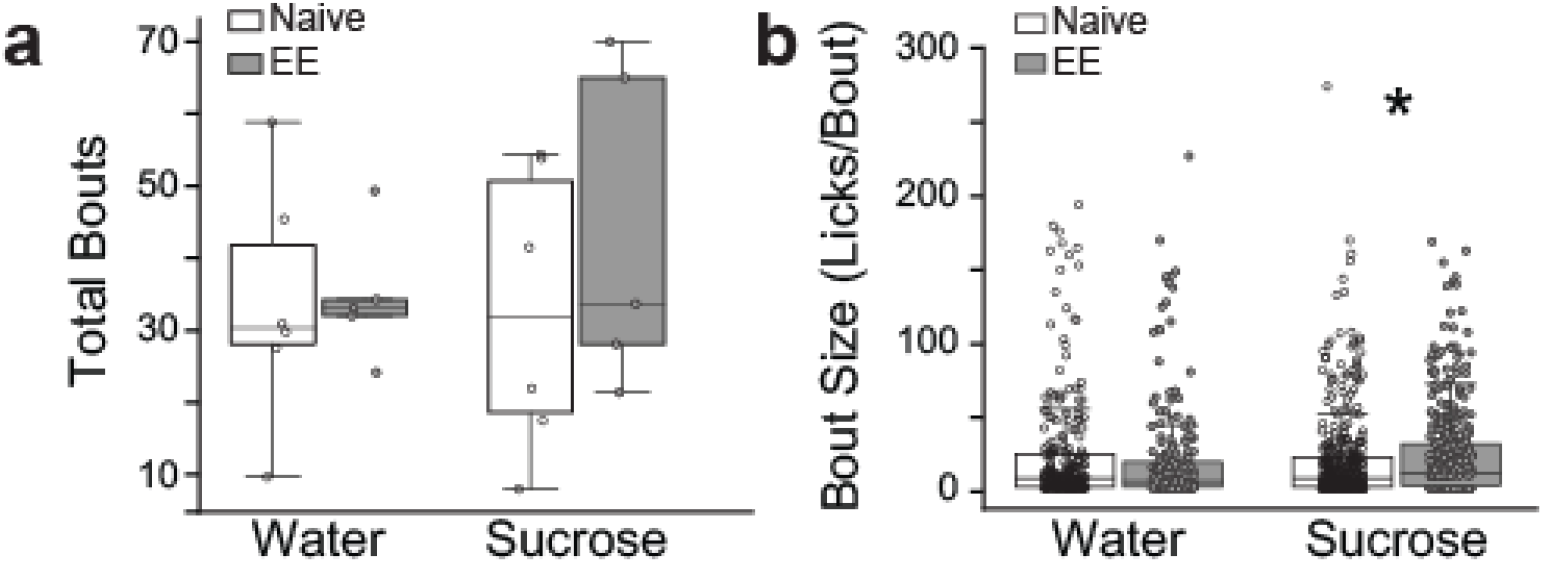
Lick bout duration is increased following EE. **a.** Total number of bouts in a 30-minute water or sucrose session were comparable between naïve (white; n=8 sessions in total from 6 mice) or EE mice (grey; n=5 mice, water p=0.4009, Mann-Whitney U test; sucrose p=0.2311, Mann-Whitney U test). **b.** Bout size, measured as licks/bout, for 30-minutes licking to water was not different between naïve and EE mice (white; naive: n=252 bouts total from 6 mice; grey; EE: n=225 bouts total from 5 mice, p=0.4828, Mann-Whitney U test). However, during 30-minutes licking to sucrose, bout size was larger following EE compared to naive (naïve: n=381 bouts total from 6 mice, EE n=434 bouts total from 5 mice, p=0.0158, Mann-Whitney U test). Asterisks indicate p ≤ 0.05.

**Supplementary Figure 3.**
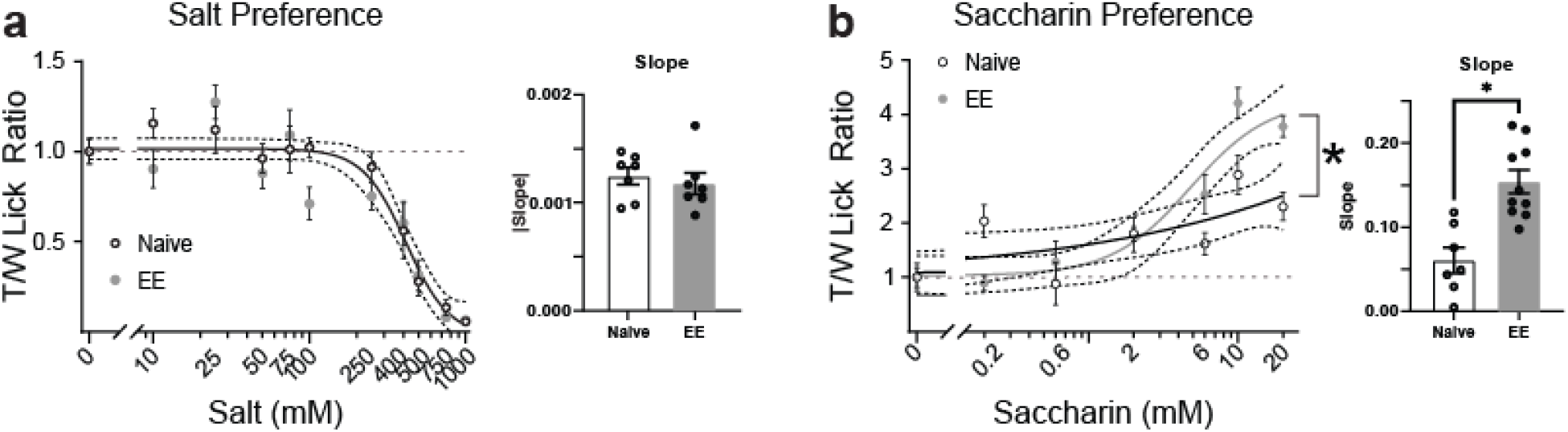
The effect of EE generalizes to saccharin, but not to salt. **a.** EE did not change salt preference, as measured with BAT (left: salt curve: naïve, black open circles n=7, EE, grey filled circles n=7 mice; F_(4,142)_=1.003, p=0.408, F test; right: absolute value of the slope: t_(12)_=0.547, p=0.595, two-tailed unpaired t-test). **b.** EE increased the preference for saccharin by shifting the preference curve (left: saccharin curve: naïve, black open circles n=7, EE, grey filled circles, n=13 mice; F_(3,110)_=7.569, p=1.195×10^-4^, F test; right: slope: t_(15)_=4.420, p=4.968×10^-4^, two-tailed unpaired t-test). Asterisks indicate p ≤ 0.05.

**Supplemental Figure 4.**
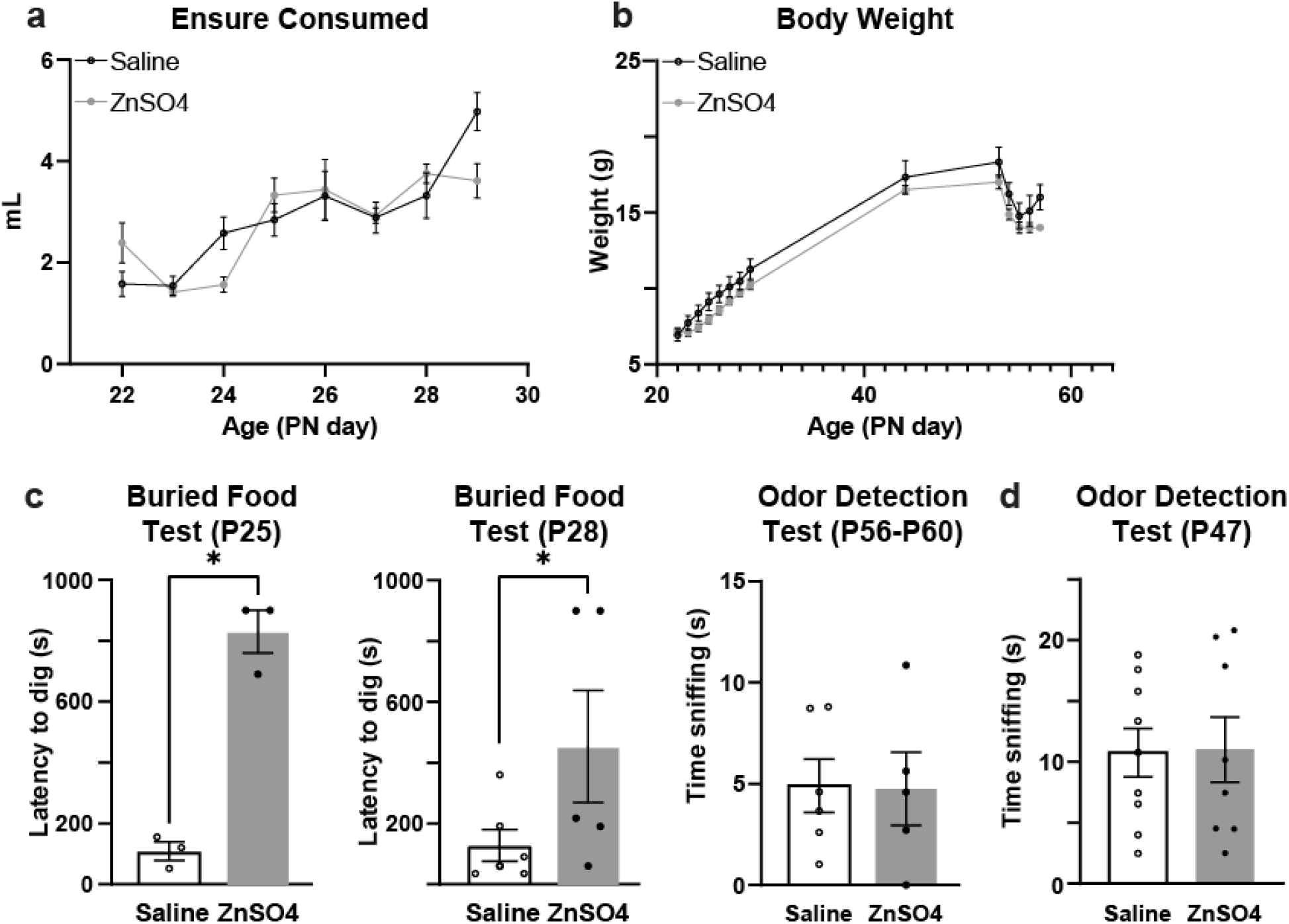
Temporary anosmia did not affect consumption, nor body weight, but impaired olfactory behavior. **a.** The amount of Ensure consumed was not different between the group that received an intranostril instillation of saline or ZnSO_4_ (grey; temporary anosmia) (black, saline n=9; grey, ZnSO_4_ n=8 mice; F_(1,15)_=0.07741, p=0.785, 2-way RM ANOVA main effect of drug). **b.** Body weight of mice treated with intranasal instillation of saline or ZnSO_4_ was not affected before, during, or after 8 days of Ensure exposure (saline, black n=9, ZnSO_4_, grey n=8 mice; F_(1,15)_=1.743, p=0.207, 2-way RM ANOVA main effect of drug). **c.** The ability to discover buried food, a behavior that relies on the sense of smell, was impaired in ZnSO_4_-treated mice compared to saline-treated mice at P25 and at P28. ZnSO_4_ resulted in an enhanced latency to dig in the proper location in a buried food test when tested at P25 (left; white, saline n=3; grey, ZnSO_4_ n=3 mice; t_(4)_=9.456, p=3.489×10^-4^, one-tailed t-test) and at P28 (middle; white, saline n=6; grey, ZnSO_4_ n=5 mice; t_(9)_=1.847, p=0.049, one-tailed t-test). The ZnSO_4_-induced anosmia was not permanent. ZnSO_4_-treated mice recovered their ability to smell in early adulthood, as assessed with a habituation/dishabituation odor detection task (right; white, saline n=6; grey, ZnSO_4_=5 mice; t_(9)_=0.06791, p=0.473, one-tailed t-test). **d.** For experimental mice later run on the BAT, no Froot Loops were used. Recovery of smell was tested at P47 using a habituation/dishabituation odor detection task to avoid the confounding effect of feeding them a sweet substance, a requirement of the buried food test. We found that recovery of smell was complete by P47. The ability to detect a novel odor was intact in a habituation/dishabituation odor task several weeks after ZnSO_4_ administration (saline n=9, ZnSO_4_ n=8 mice; t_(15)_=0.110, p=0.457; unpaired one-tailed t-test). Asterisks indicate p ≤ 0.05.

**Supplementary Figure 5.**
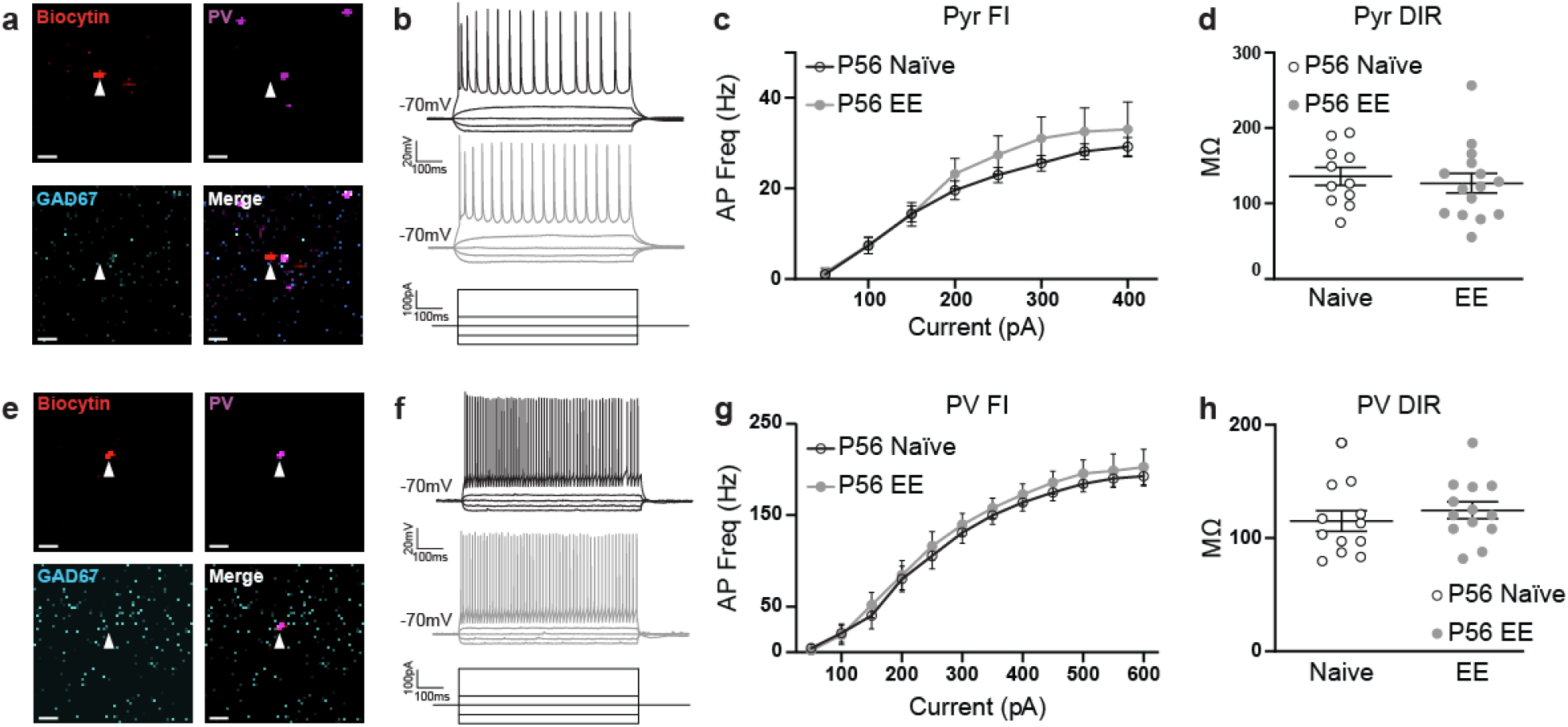
EE does not alter the intrinsic excitability of PYR or PV^+^ neurons. **a, e.** Sample recorded PYR neuron (a) and PV^+^ neuron (e) with biocytin fill in red (top left), tdTomato signal in magenta (top right), GAD67 immunostain in cyan (bottom left), and the merged image (bottom right). Images show a single focal plane. White arrowhead points to location of the recorded neuron. PYR neurons were confirmed by the absence of colocalization with GAD67, while PV^+^ neurons colocalize with GAD67. **b, f.** Sample traces of PYR neuron (b) and PV^+^ neuron (f) from naïve (black, top) or EE (grey, middle) mice recorded in current clamp, holding neurons at −70mV. Bottom diagram shows sample current inputs for the corresponding voltage traces. **c, g.** Input/output function for GC PYR neurons (c) and PV^+^ neurons (g) recorded in slices from naïve (black) and EE (grey) mice. There were no differences in input/output function in either neuron type following EE (naïve n=11 cells from 5 mice, EE n=15 cells from 5 mice, F_(1,24)_=0.539, p=0.470, 2-way RM ANOVA, main effect of exposure). **d, h.** The dynamic input resistance of GC PYR neurons (d) and PV^+^ neurons (h) recorded in slices from naïve and EE mice was not significantly different (naïve n=11 cells from 5 mice, EE n=15 cells from 5 mice, t_(24)_=0.501, p=0.621, two-tailed t-test).

**Supplementary Figure 6.**
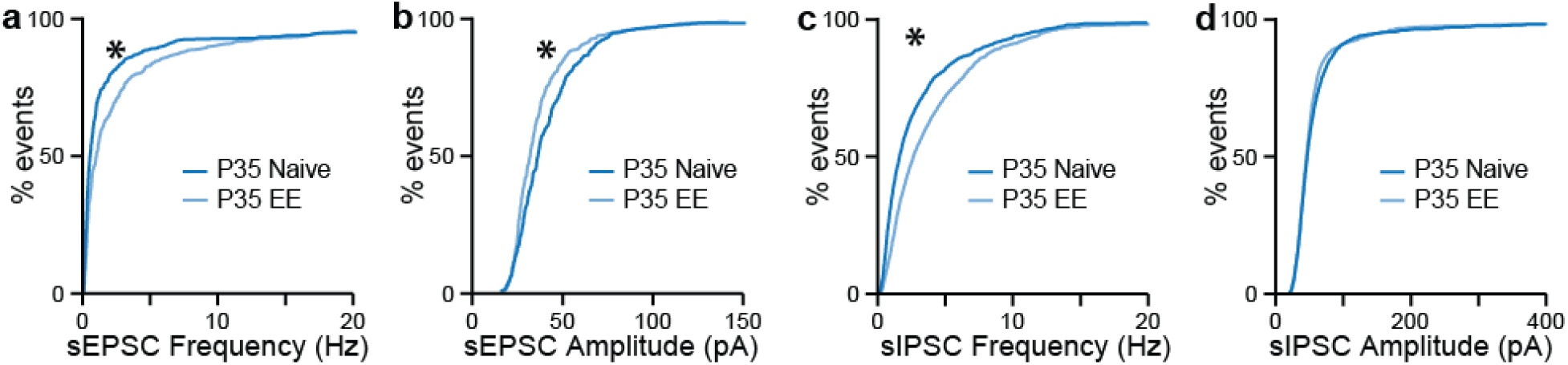
EE induces transient changes sEPSCs and long lasting increase in sIPSCs. To determine whether the change in inhibitory synaptic transmission was a long-term effect of EE or a change that emerges soon after exposure, we performed voltage clamp recordings in naïve and EE wild-type mice at P35, only a few days after the end of exposure. **a.** Cumulative distributions of sEPSCs frequency (sEPSCs frequency, naïve (dark blue) n=10 cells from 5 mice; EE (light blue) n=9 cells from 6 mice; D=0.191, p=1.192×10^-5^, K-S test). **b.** Cumulative distribution of sEPSCs amplitude (sEPSPc amplitude, naïve (dark blue) n=10 cells from 5 mice; EE (light blue) n=9 cells from 6 mice; D=0.171, p=1.172×10^-4^, K-S test). **c.** Cumulative histograms showing increased sIPSCs frequency (sIPSCs frequency, naïve (dark blue) n=18 cells from 9 mice; EE (light blue) n=14 cells from 6 mice, D=0.1873, p<1.0×10^-14^, K-S test). **d.** Cumulative distribution of sIPSC amplitude (naïve n=18 cells from 9 mice, EE n=14 cells from 6 mice, D=0.0514, p=0.031, K-S test; threshold for significance p=0.01). Together, these results indicate that EE induced a transient early modulation of excitatory synaptic transmission, and a persisten increase in inhibitory synaptic transmission. Asterisks indicate p ≤ 0.05.

**Supplementary Figure 7.**
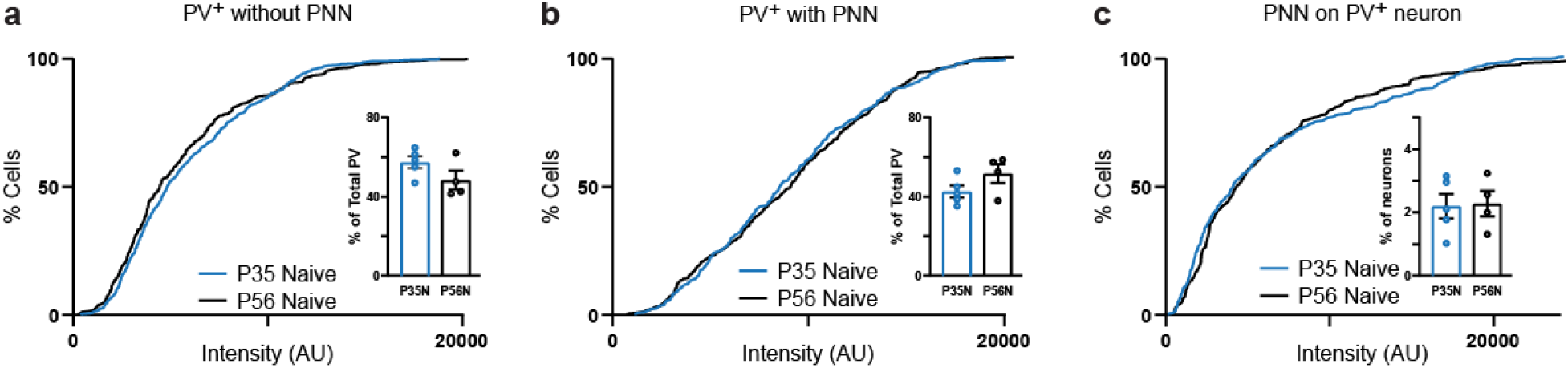
Stable expression of PV and association with PNNs between P35 and P56 in control animals. **a.** The PV fluorescence intensity signal of PV^+^ neurons without a PNN was stable from P35 to P56 (P35 n=503 neurons from 5 mice, P56 n=317 neurons from 4 mice; D=0.07039, p=0.290). Inset: the percent of PV^+^ neurons without a PNN was not changed from P35 to P56 (P35 n=5, P56 n=4 mice; t_(7)_=1.654, p=0.142, two-tailed unpaired t-test). **b.** The PV fluorescence intensity signal of PV^+^ neurons with a PNN was stable from P35 to P56 (P35 n=382 neurons from 5 mice, P56 n=343 neurons from 4 mice; D=0.04685, p=0.822, K-S test). Inset: the percent of PV^+^ neurons with a PNN was not changed from P35 to P56 (inverse of data shown in the inset of panel A: P35 n=5, P56 n=4 mice; t_(7)_=1.654, p=0.142, two-tailed unpaired t-test). **c.** WFA fluorescence intensity was stable from P35 to P56 (P35 n=379 PNNs from 5 mice, P56 n=344 PNNs from 4 mice; D=0.0865, p=0.134, K-S test). Inset: the percent of PNNs out of total neurons was not changed from P35 to P56 (P35 n=5, P56 n=4 mice; t_(7)_=0.1387, p=0.894, two-tailed unpaired t-test).

